# Mechanisms controlling the deposition and dynamics of histone variant H2BE

**DOI:** 10.64898/2026.04.20.719605

**Authors:** Annabel K Sangree, Emily R Feierman, Roxanne Perez-Tremble, Erica Korb

## Abstract

Histone variants shape chromatin structure and gene regulation and their deposition and localization in chromatin are tightly controlled. H2BE is the only known widely expressed mammalian H2B variant and localizes to transcription start sites to control chromatin accessibility and cognitive function. However, the mechanisms governing H2B variant incorporation into chromatin remain unclear. Here, we define the regulatory framework controlling H2BE incorporation, localization, and eviction in neuronal chromatin. We identify the BAF remodeling complex and the transcription factor SP1 as key drivers of H2BE deposition at specific genomic loci. We further show that FACT maintains H2BE enrichment at transcription start sites by preventing its distribution into gene bodies, while the histone chaperone NAP1L4 mediates H2BE eviction from chromatin. Comparative analysis with H2A.Z reveals that BAF, SP1 and NAP1L4 exert H2BE-specific functions, while FACT functions more broadly across histone variants. Finally, we define the H2BE-dependent transcriptional effects of it chaperones. Together, these findings uncover the first chaperones governing H2B variant incorporation and define a highly complex mechanism responsible for H2BE regulation in chromatin.

**Graphical abstract:** 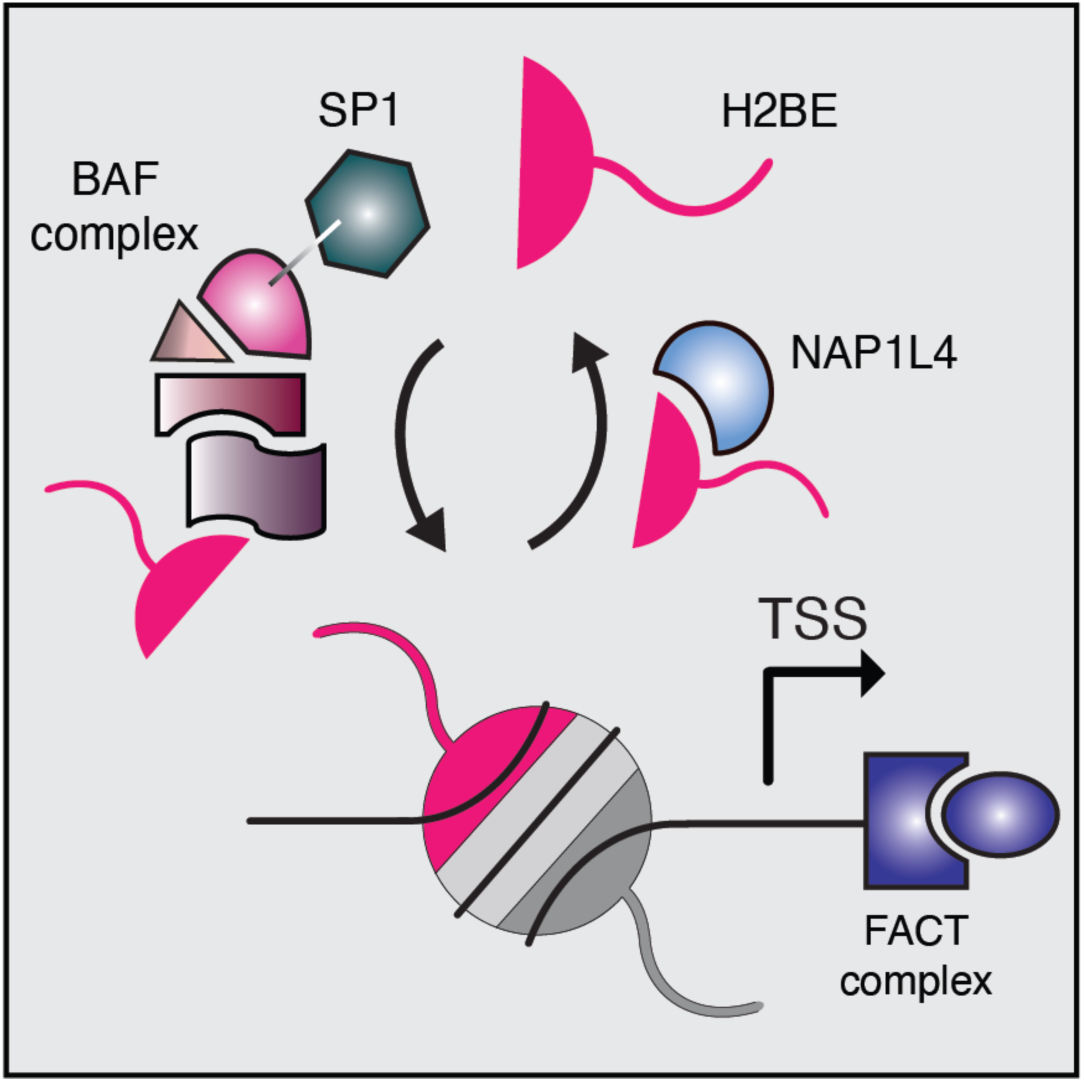

**Highlights:**

- The BAF complex controls H2BE incorporation.
- SP1 promotes locus-specific incorporation of H2BE.
- FACT prevents H2BE accumulation in genic regions.
- NAP1L4 mediates eviction of H2BE from chromatin.

## Introduction

Histone proteins are fundamental regulators of genome organization that control chromatin architecture and transcriptional output. Canonical histones are broadly expressed and encoded within large multigene clusters containing 10–20 nearly identical copies. By contrast, histone variants are typically encoded by only one or two genes and contain specific amino acid substitutions compared to their canonical counterparts. This allows histone variants to create distinct chromatin environments that affect transcription^1^. Unlike canonical histones, variants are also generated through replication-independent mechanisms and typically accumulate in non-dividing cells. In post-mitotic cells such as neurons, which lack replication–coupled chromatin assembly, genome stability and transcriptional regulation rely heavily on the incorporation of histone variants^1–3^.

Histone variants are incorporated by specific chaperones or ATP-dependent chromatin remodelers^4,5^. These chaperone proteins facilitate the assembly, disassembly, and exchange of histone dimers within the nucleosome core. Variants like H3.3 have specific chaperones that recognize sequence motifs present only in the variant form^6,7^. The deposition of other variants, such as H2A.Z, is facilitated by the ATP-dependent chromatin remodeling SRCAP complex^8,9^. Others, like MacroH2A, are regulated in part by chaperone complexes that act on canonical histones and are also deposited by variant-specific chaperones^5^. Disruption of these and other histone chaperones can lead to numerous neurodevelopmental disorders^10–12^.

Unlike other histone proteins, H2B variants remain highly understudied. H2BE is the only known mammalian H2B variant expressed in many tissues and like many histone variants, it plays critical roles in the brain^13–15^. H2BE was initially identified as a regulator of neuronal longevity in the mouse olfactory system^16^. Recent work demonstrated that H2BE is expressed in numerous tissues and is highly abundant in multiple brain regions^17^, and particularly enriched in neurons and astrocytes^18^. H2BE localizes specifically to transcription start sites (TSSs) of actively transcribed genes and is essential for neuronal transcription, homeostatic scaling, and long-term memory^17,19^. In addition to its role in neurons, H2BE promotes gene expression in astrocytes and contributes to transcriptional regulation in the aging brain^18^. However, despite H2BE’s broad importance in transcriptional regulation and brain function, the mechanisms that regulate its localization within chromatin, including its targeted deposition and eviction, remains largely unknown.

The unique localization of H2BE to the TSS suggests the existence of dedicated chaperone proteins or regulatory mechanisms that distinguish it from canonical H2B. Canonical H2B is deposited throughout the genome by well-characterized chaperones, including Nucleosome Assembly Protein 1 (NAP-1)^20–22^ and Facilitates Chromatin Transcription (FACT)^23,24^. H2BE differs from canonical H2B by only five amino acids distributed across the protein^16^, rather than a discrete targeting motif typically observed in other variants. These subtle sequence differences may be insufficient to allow for selective regulation by dedicated chaperones. Alternatively, H2BE incorporation may rely on additional factors beyond classical histone chaperones to achieve its restricted localization such as through association with multiple chromatin regulators that act at distinct genomic regions. However, no chaperone proteins or other regulatory partners, whether shared with canonical H2B or unique to the variant, have been identified for H2BE.

Here, we define the first model governing H2BE incorporation, eviction, and localization in neuronal chromatin by applying biochemical and sequencing approaches. We find that subunits of the ATP-dependent remodeling BAF complex associate with H2BE, and together with the transcription factor SP1, promote its incorporation into chromatin at specific loci. Conversely, NAP1L4, a member of the NAP-1 family, mediates H2BE eviction from chromatin, providing an additional layer of regulatory control. Further, FACT maintains H2BE’s positioning at the TSS during transcription by restricting it from gene bodies. Parallel CUT&Tag analysis of H2A.Z demonstrates that BAF, SP1 and NAP1L4 exert largely H2BE-specific functions, while FACT has similar effects across histone variants. Finally, we define the effects of NAP1L and BAF on transcription that are dependent on H2BE. Collectively, these findings establish a regulatory framework for H2BE dynamics and reveal how this histone variant contributes to neuronal chromatin organization and transcriptional responses. Further, they define the first known mechanisms controlling the incorporation and eviction of a mammalian H2B variant protein.

## Results

### Identification of H2BE binding partners

While recent findings demonstrate the importance of H2BE in multiple cell types^16–19^, the regulation of its genomic incorporation, localization, and eviction remains unknown. Here, we sought to identify the mechanisms that control H2BE in chromatin by first identifying the proteins with which it associates. We performed co-immunoprecipitations (CoIPs) in primary cortical neurons from E16.5 embryos following 12 days in culture using an antibody specific for H2BE, which we previously validated^17^. In parallel, we performed CoIPs targeting canonical H2B and then used mass spectrometry (MS) to identify associated proteins for H2B and H2BE (Figure 1A, Figure S1A). We first validated successful and specific CoIPs, focusing on the I39 residue on H2BE (V39 in H2B), which allowed for the efficient differentiation between peptides (Figure S1B). As anticipated, we found enriched H2BE in the H2BE CoIP and H2B in the H2B CoIP (Figure S1A). Interestingly, we could not detect H2B in the H2BE CoIP condition or vice versa, despite each associating similarly with other canonical histones (Figure 1B, C; S1B). This suggests that nucleosomes may generally be composed of either variant H2BE *or* canonical H2B, with mixed nucleosomes being less common.

**Figure 1.**
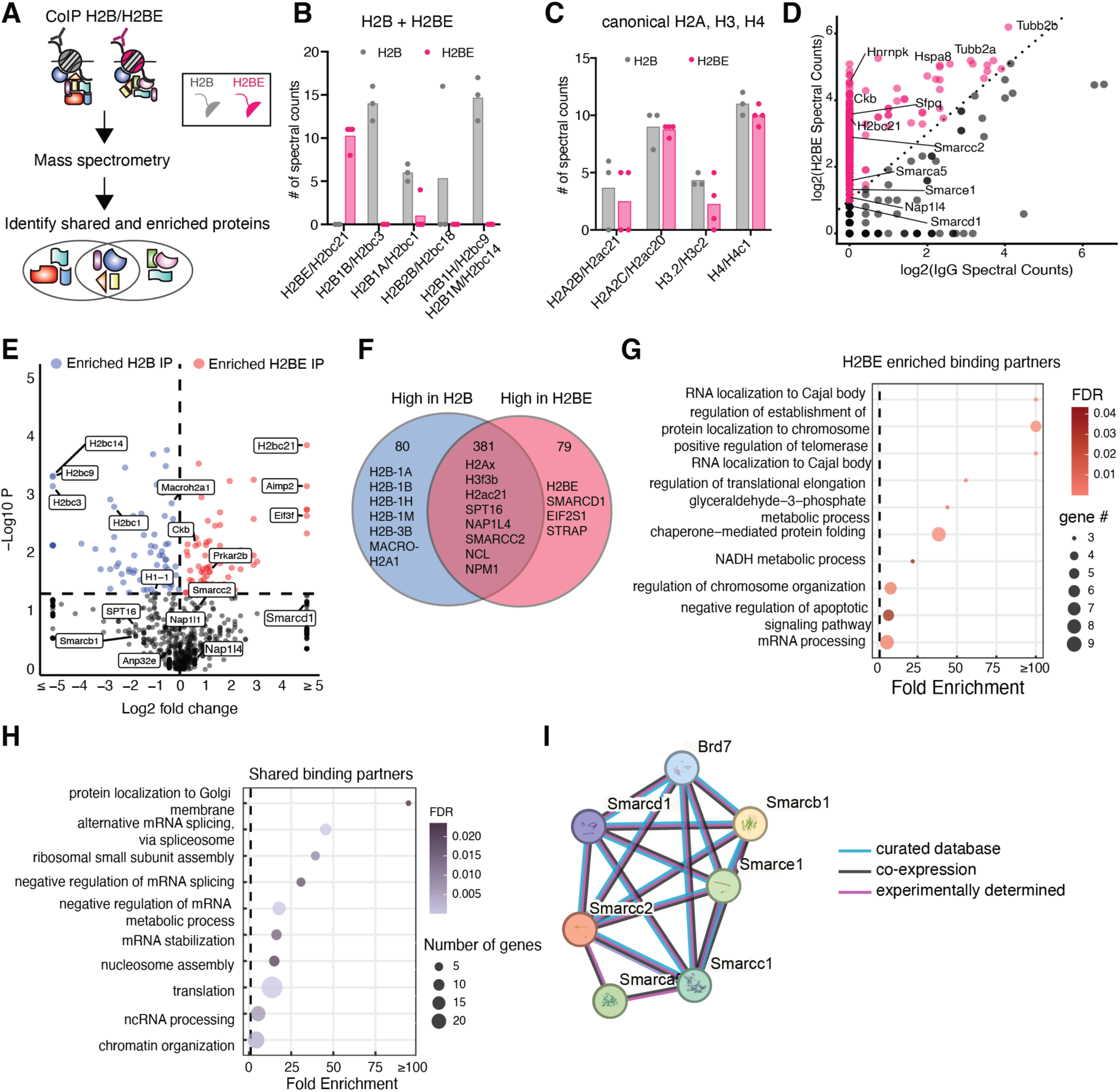
Identification of H2BE binding partners via mass spectrometry. A) Experimental setup. Primary cortical neurons from E16.5 embryos were cultured for 12 days, after which Coimmunoprecipitations (CoIPs) for H2B and H2BE were performed. The resulting products were analyzed using mass spectrometry. B) Number of spectral counts of either H2BE (pink) or H2B (grey) with either H2BE or H2B histone proteins. C) Same as B, but for canonical H2A, H3 and H4 histones. D) Scatter plot of log2(H2BE spectral counts) vs log2(IgG spectral counts). The dashed line represents a cutoff of 2-fold higher spectral counts in H2BE IP vs IgG. Points above this threshold are colored in pink. E) Volcano plot of interacting partners colored by enrichment in H2B (blue), H2BE (red), or shared (grey). F) Venn diagram of results in D by proteins that were high in H2B, high in H2BE, or shared. G) GO plot of H2BE-enriched binding partners. H) GO plot of shared binding partners. I) STRING module of BAF complex members identified as shared or H2BE-enriched binding partners.

We next examined the enrichment of H2BE binding partners compared to an IgG control and identified several known chaperones H2B chaperones and chromatin remodelers, including multiple members of the BAF complex, the NAP1 family, and FACT complex member SPT16 (Figure 1D). These data align with previous work done using FLAG-H2BE overexpression in the mouse main olfactory epithelium, in which BAF and FACT complex proteins were identified as H2BE binding partners^16^. Comparing H2B and H2BE associations, we found that canonical H2B is preferentially associated with MacroH2A1/2 and several H1 linker proteins compared to H2BE (Figure S1C, D). As MacroH2A and H1 are typically associated with closed chromatin^25–28^, this is consistent with the role of H2BE in promoting open chromatin and localizing to active TSS regions^17^. Also fitting with H2BE’s function in promoting transcription, we found that signaling proteins linked to transcriptional activation like STRAP and CKB are more highly associated with H2BE (Figure 1D-F), while many heterochromatic proteins are enriched in H2B-associated proteins (Figure S1E). Together, these findings support the established role of H2BE in promoting chromatin accessibility to maintain transcription of highly expressed genes^17^.

Finally, we examined both H2BE-enriched associated proteins for known chaperone proteins. H2BE is associated with several established H2B chaperones and chromatin remodelers, including members of the BAF complex, the NAP1 family, and FACT complex member SPT16 (Figure 1E, H). Notably, both H2B and H2BE associate with these putative chaperones. This suggests that, unlike many other histone variants, because H2BE is highly similar to canonical H2B and lacks a cluster of unique amino acids that could recruit a specific chaperone protein, it may instead share chaperone proteins with H2B. Together, these analyses identified proteins that preferentially associate with H2B or H2BE and identify several known chaperone proteins that associate with both H2B and H2BE.

### The BAF complex promotes the incorporation of H2BE into chromatin

Among the many candidate chaperone proteins identified, we observed a notable enrichment of many members of the BAF complex as either shared or even H2BE-specific binding partners by MS (Figure 1I). The BAF complex is an ATP-dependent chromatin remodeler that can initiate nucleosome remodeling, including through contact with the H2A-H2B dimer^29^. Other histone variants such as H2A.Z are deposited by similar ATP-dependent remodeling complexes^8,30^. We hypothesized that H2BE incorporation may likewise be mediated by an ATP-dependent remodeling mechanism and incorporated as the BAF complex acts on nucleosomes. To test this, we first confirmed that H2BE associates with subunits of the BAF complex in N2A cells by Co-IP of FLAG-tagged H2BE and probing for BAF subunits SMARCC2, SMARCB1, and SMARCD1 (Figure 2A). Next, to disrupt BAF complex activity, we used the compound *BRD-K98645985* (BAFi), which inhibits the binding of ARID1A-containing BAF (cBAF) complexes to chromatin^31–33^. We treated primary cortical neurons from E16.5 embryos with either DMSO or 10 μM of BAFi for 6 and 24 hours and assessed H2BE expression in protein lysates. H2BE expression levels were stable across DMSO and BAFi treatments (Figure S2A), demonstrating that inhibiting the BAF complex does not affect global H2BE levels.

**Figure 2.**
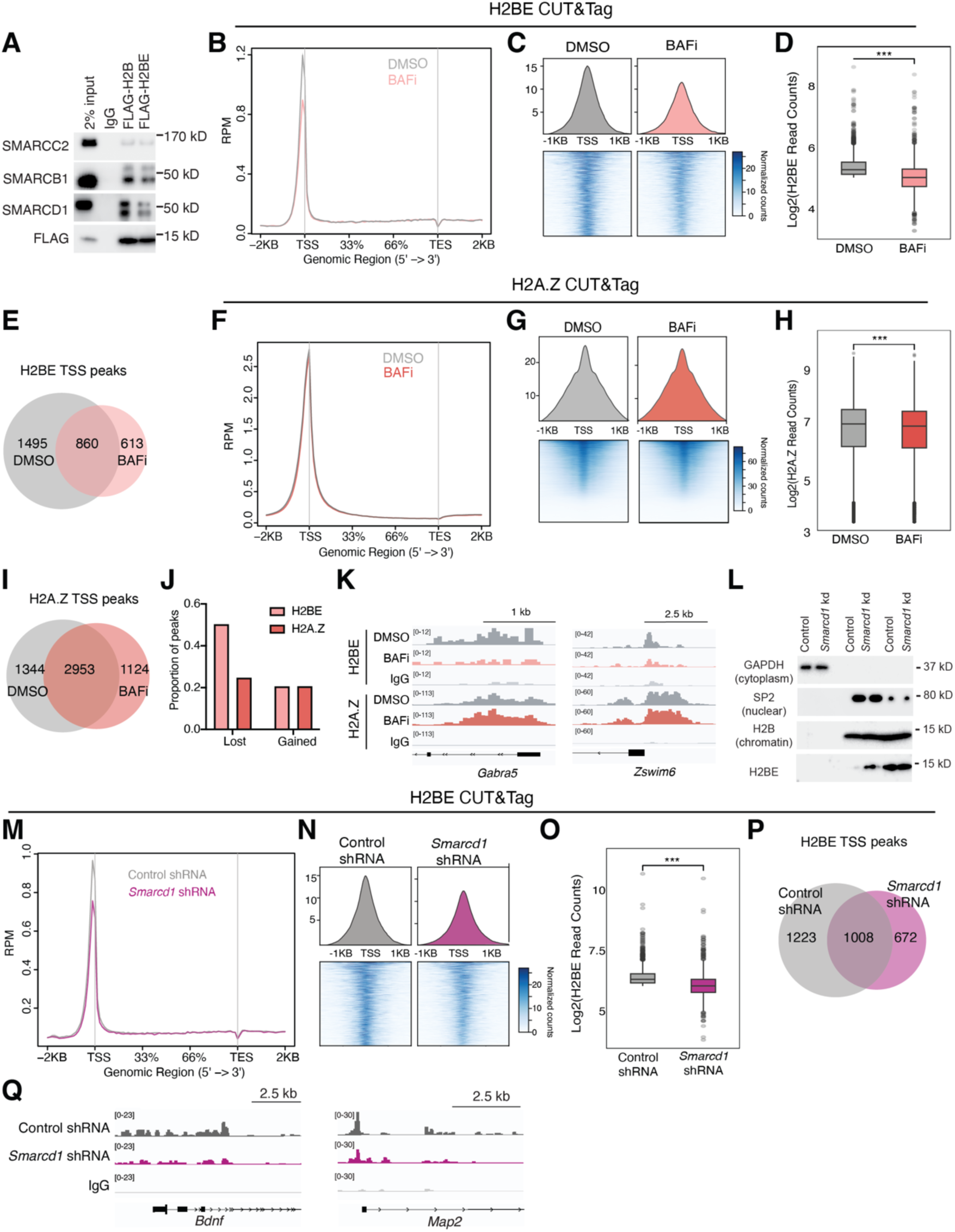
The BAF complex regulates H2BE in chromatin. A) CoIP for FLAG-tagged H2B and H2BE and BAF subunits SMARCC2, SMARCB1, and SMARCD1 in N2A cells. B) Metaplot of H2BE CUT&Tag profiling in WT primary cortical neurons treated with DMSO (grey) or BAFi (pink). The plot shows read counts per million mapped reads (RPM) at ± 2 kb from the TSS of genes with a high confidence H2BE peak (n = 2 biological replicates per treatment). C) Heatmap and metaplot of H2BE signal +/- 500bp of the TSSs of genes with a high confidence H2BE peak (n = 2357 genes). D) Normalized H2BE CUT&Tag read counts of high confidence H2BE peaks +/- 1KB of the TSS. n = 2357 genes. (unpaired t test; ***, p < 0.001). E) Overlap of H2BE TSS peaks called by SEACR between DMSO and BAFi (peak score = 12). F) Metaplot of H2A.Z CUT&Tag profiling in WT primary cortical neurons treated with DMSO (grey) or BAFi (orange). n = 3 biological replicates per treatment. G) Heatmap and metaplot of H2A.Z signal +/- 1KB of the TSS across all genes. H) Normalized H2A.Z CUT&Tag read counts at H2A.Z peak sites with ≥ 10 reads detected at +/- 500bp of the TSS (unpaired t test; ***, p < 0.001). I) Overlap of H2A.Z TSS peaks called by SEACR between DMSO and BAFi (peak score = 50). J) Proportion of lost and gained peaks for H2BE (pink) and H2A.Z (orange) CUT&Tag with BAFi. K) CUT&Tag gene tracks for representative genes, with bigwigs merged across replicates. H2BE is shown at the top, followed by H2A.Z. L) Subcellular fractionation of N2A cells transduced with either Control or *Smarcd1* shRNAs. GAPDH marks the cytoplasmic fraction, SP2 the nuclear, and H2B the chromatin-bound fraction. M) Metaplot of H2BE CUT&Tag profiling in WT primary cortical neurons transduced with a Control shRNA (grey) or *Smarcd1* targeting shRNA (pink) n = 3 biological replicates per treatment. The plot shows read counts per million mapped reads (RPM) at ± 2 kb from the TSS of genes with a high confidence H2BE peak. N) Heatmap and metaplot of H2BE signal +/- 1KB of the TSSs of genes with a high confidence H2BE peak (n = 2798 genes). O) Normalized H2BE CUT&Tag read counts at H2BE peak sites with ≥ 10 reads detected +/- 500bp of the TSS (unpaired t test; ***, p < 0.001). P) Overlap of H2BE TSS peaks called by SEACR between Control and *Smarcd1* depletion (peak score cutoff = 15). Q) CUT&Tag gene tracks for representative genes. Bigwigs were generated from individual replicates and then merged.

To identify genomic localization changes induced by BAF complex inhibition, we used cleavage under targets and tagmentation (CUT&Tag) with sequencing^34^. We treated primary cortical neurons with either DMSO or BAFi for 24 hours before collecting neurons for CUT&Tag. In alignment our previous work^17^, H2BE was highly enriched at the TSS (Figure 2B, S2B). However, BAFi resulted in significantly less H2BE at the TSS compared to control both genome-wide and at genes with a high-confidence H2BE peak (Figure 2B-D, S2B-D). Peak calling revealed 1,495 H2BE peaks lost and 613 peaks gained upon BAFi treatment (Figure 2E). Lost H2BE peaks were at the TSSs of genes related to cellular and protein localization, including many genes specifically important for neuronal function (Figure S2E, 2F). BAFi decreased H2BE at genes ubiquitously, regardless of expression level (Figure S2F). Together, these results indicate that BAF activity promotes H2BE incorporation at the TSSs, including at genes important for neuronal function.

To assess the specificity of the BAF complex in promoting H2BE incorporation, we performed CUT&Tag for another histone variant, H2A.Z, in neurons treated with BAFi. We detected only a subtle decrease in H2A.Z deposition at the TSS with BAFi compared to DMSO (Figure 2F-H). Notably, while H2A.Z had more than double the number of H2BE peaks at baseline, BAFi induced a greater loss of H2BE peaks than H2A.Z peaks (Figure 2I–K, S2G-I), indicating that this complex preferentially promotes H2BE deposition.

### Smarcd1 (BAF60A) depletion recapitulates BAF inhibition

To validate the role of the BAF complex in H2BE regulation using a genetic approach, we selected SMARCD1 (BAF60A) as a target. SMARCD1 is the only SMARCD core-subunit present in the MS data, it interacts with H2BE (Figure 2A) and is incorporated into all BAF complex assemblies^35^. Importantly, loss of SMARCD paralogs disrupts BAF complex assembly and prevents incorporation of both ARID and ATPase subunits^35,36^, making SMARCD1 an effective point of genetic inhibition for the entire complex. Notably, loss-of-function mutations in SMARCD1 cause a neurodevelopmental disorder^12^, highlighting the importance of SMARCD1 in the brain.

We designed shRNAs targeting *Smarcd1* and tested knockdown efficiency in both N2A cells and neurons. In both cell types, we achieved a robust knockdown of the *Smarcd1* transcript (Figure S2J) and partial knockdown at the protein level (Figure S2K). As with general BAF inhibition, we did not observe global changes in H2BE expression upon *Smarcd1* depletion (Figure S2K). Next, we performed a subcellular fractionation, isolating the cytoplasmic, nuclear, and chromatin fractions of N2A cells, and found that *Smarcd1* depletion increased H2BE in the nuclear fraction (Figure 2L). These results indicate that the BAF complex promotes H2BE chromatin incorporation, as loss of *Smarcd1* prevents full H2BE incorporation into chromatin.

Finally, we assessed the consequences of reduced H2BE incorporation with *Smarcd1* depletion using CUT&Tag in primary cortical neurons. Consistent with previous observations using BAFi, *Smarcd1* depletion resulted in significantly less H2BE at the TSS both genome-wide and at high-confidence H2BE peaks (Figure 2M-P, S2L-N). *Smarcd1* depletion also resulted in a significant loss of H2BE at the TSS of genes related to RNA processing, metabolic processing, and localization (Figure 2Q, S2O), and reduction occurred independently of gene expression level (Figure S2P). Further, we assessed the overlap between the BAFi and *Smarcd1* depletion datasets and observed a high concordance between datasets (Pearson r = 0.8, Figure S2Q). Together, these orthogonal approaches indicate that the BAF complex incorporates H2BE into chromatin.

### SP1 restricts BAF-mediated H2BE deposition

Given that the BAF complex acts widely across the genome, yet only a portion of genes contain H2BE, we next sought to identify how H2BE is directed to its target promoters. Because H2BE deposition positively correlates with transcription^17^, we hypothesized that a sequence-specific transcription factor might bridge this gap. Motif analysis of promoters of genes with H2BE^17^ identified numerous SP transcription factor family members with the SP1 consensus sequence as the most enriched hit (Figure 3A). While SP1 motifs are highly abundant throughout the genome, SP1 is known to associate with members of the canonical BAF complex^37,38^, and thus the two acting together could jointly dictate where H2BE is deposited. We first validated the association of SP1 with both H2BE and SMARCD1 in N2As (Figure 3B, S3A). Next, using BAF complex protein BRG1 (SMARCA4) CUT&RUN data^39^, we quantified H2BE occupancy under baseline conditions at the TSSs of genes that have an SP1 motif but no BRG1 peak; a BRG1 peak but no SP1 motif, or those that had both a BRG1 peak and SP1 motif (Figure 3C). Supporting our hypothesis, we found that H2BE was most abundant at genes with both a BRG1 peak and an SP1 motif.

**Figure 3.**
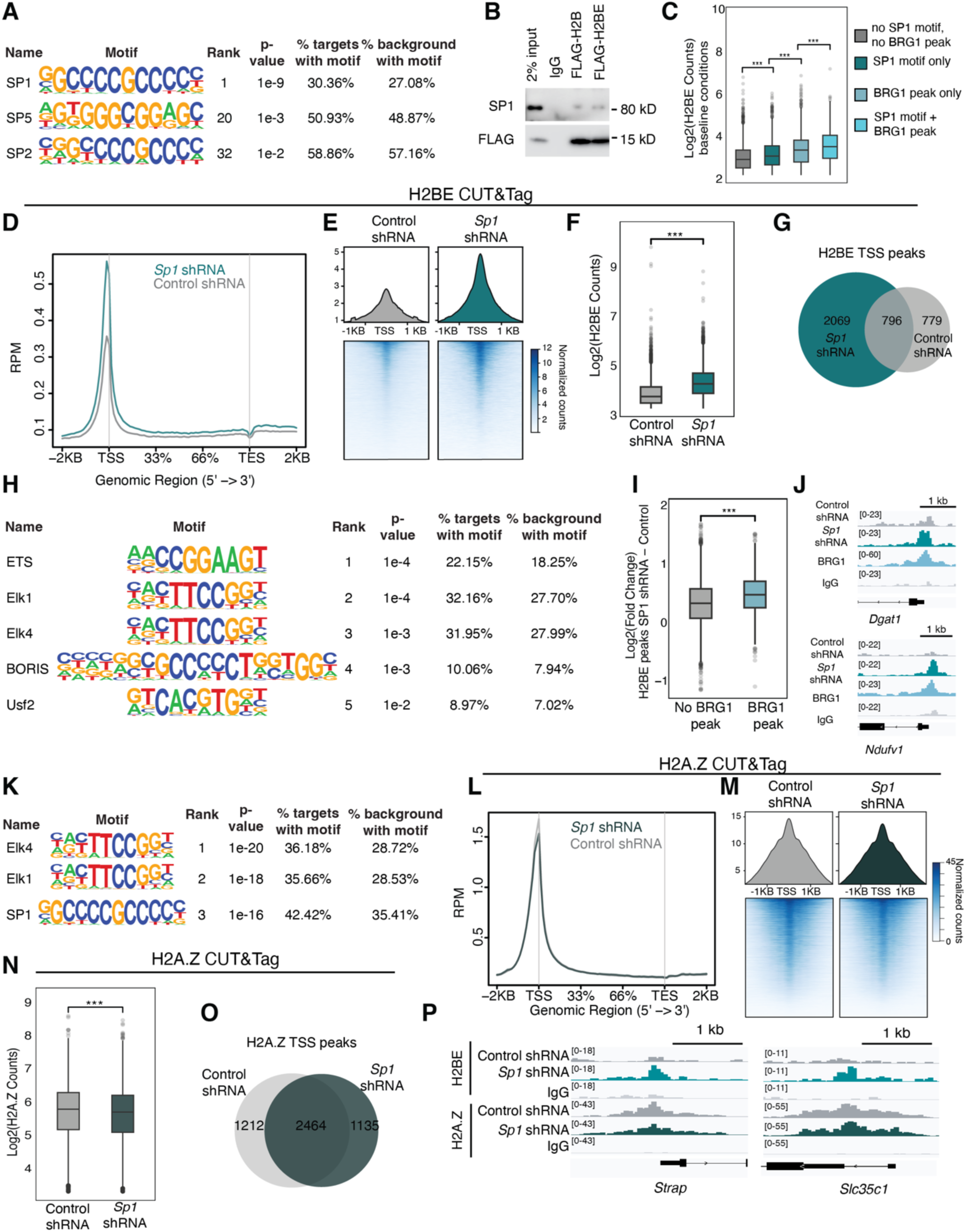
SP1 restricts BAF-mediated H2BE deposition. A) Motif analysis of genes with H2BE at their TSS in WT E16.5 primary cortical neurons (data from^17^) identifies enrichment of members of the SP transcription factor family. B) CoIP for FLAG-tagged H2BE and SP1 in N2A cells. C) Normalized H2BE read counts at baseline binned by the presence of a BRG1 peak (data from^39^) and/or an SP1 motif. Significance was calculated using a one-way ANOVA, followed by a post hoc Tukey’s HSD test. *** p, 0.001. D) Metaplot of H2BE CUT&Tag profiling in WT primary cortical neurons transduced with a Control (grey) or *Sp1*-targeting (green) shRNA. Plot shows read counts per million mapped reads (RPM) at ± 2 kb from the transcription start site (TSS) (*n* = 3 biological replicates per condition). E) Heatmap and metaplot depicting H2BE at +/- 1KB from the TSS. F) Normalized H2BE CUT&Tag read counts at H2BE peak sites with ≥ 10 reads detected +/- 500bp from the TSS (unpaired t test, *** p <0.001). G). Overlap of H2BE TSS peaks called by SEACR between Control and *Sp1* shRNAs (peak score = 7). H) Motif analysis of genes that gained H2BE at their TSS with *Sp1* depletion. Quantification of log2-fold change (*Sp1* depletion – Control) of H2BE at genes without (grey) or with (light blue) a BRG1 peak (unpaired t test, *** p <0.001). J) Representative gene tracks of genes that gain H2BE at their TSS with *Sp1* depletion and have a BRG1 peak at baseline. K) Motif analysis of genes that with H2A.Z at their TSS under baseline conditions. L) Metaplot of H2A.Z CUT&Tag profiling in WT primary cortical neurons transduced with a Control (grey) or *Sp1*-targeting (green) shRNA. M) Heatmap and metaplot of H2A.Z signal +/- 1KB of the TSS across all genes. N) Normalized H2A.Z CUT&Tag read counts at H2A.Z peak sites with ≥ 10 reads detected at +/- 500bp of the TSS (unpaired t test, *** p <0.001). O) Overlap of H2A.Z TSS peaks called by SEACR between Control and *Sp1* depletion (peak score = 25). P) CUT&Tag gene tracks for representative genes, with bigwigs merged across replicates. H2BE is shown at the top, followed by H2A.Z.

Next, we depleted *Sp1* to define its effects on H2BE and confirmed robust knockdown on its mRNA and protein (Figure S3B-C). To identify genomic localization changes induced by *Sp1* depletion, we used CUT&Tag and found genome-wide increased deposition of H2BE, that was most notable at the TSS (Figure 3D-F). Neurons gained more than 2000 H2BE peaks upon *Sp1* depletion compared to control (Figure 3G). Motif analysis of the genes that gained H2BE with *Sp1* depletion revealed no enrichment for SP1 motifs (Figure 3H). Instead, it identified transcription factors like Elk1/4 and ETS1, which are also known to interact with the BAF complex^40,41^, suggesting aberrant H2BE deposition to other sites at which BAF can act. To further interrogate this relationship, we calculated the fold change of H2BE peaks with *Sp1* depletion at the TSS of genes with or without a BRG1 peak and observed a significantly greater gain at genes with a BRG1 peak than those without (Figure 3I-J). Together, these findings support a model in which SP1 normally restricts BAF-mediated H2BE deposition. In the absence of SP1, BAF appears to deposit H2BE more broadly across the genome, leading to non-specific expansion of H2BE occupancy.

To determine whether the changes in H2BE localization observed following *Sp1* depletion could be explained by secondary effects on transcription rather than by a direct role for SP1 in specifying H2BE deposition, we inhibited transcription using Flavopiridol^42^. Primary cortical neurons were treated with Flavopiridol for 24 hours and subjected to H2BE CUT&Tag. In contrast to the widespread increase in H2BE following *Sp1* depletion, transcriptional inhibition produced a modest increase in H2BE signal at TSSs relative to DMSO controls (Figure S3E - H). This outcome is consistent with previous reports showing that histone variant turnover is tightly coupled to active transcription, such that blocking transcription reduces nucleosome exchange and stabilizes existing histone variant incorporation^43–46^. We next assessed the overlap of genes gaining H2BE peaks following transcriptional inhibition with Flavopiridol versus *Sp1* depletion. Whereas Flavopiridol treatment resulted in relatively limited H2BE redistribution, *Sp1* depletion led to a much broader increase in H2BE across the genome (Figure S3I–J). This divergence indicates that the altered H2BE landscape observed upon SP1 loss is unlikely to be explained by reduced transcriptional activity alone.

Finally, to understand the specificity of SP1’s participation in dictating H2BE’s localization, we performed a motif analysis of genes with H2A.Z peaks (Figure 3K). SP1 was identified but ranked third rather than the top hit as found in H2BE while other top hits were those that only appeared in H2BE peaks following SP1 loss (such as ELK1 and ELK4). We next performed CUT&Tag for H2A.Z using the same Control and *Sp1* shRNAs. Contrary to what we observed with H2BE, *Sp1* depletion caused a subtle *decrease* in H2A.Z at the TSS (Figure 3L-P), supporting a model in which SP1 selectively restricts BAF-mediated deposition of H2BE. Together, these results demonstrate that SP1 selectively regulates H2BE localization, rather than broadly controlling histone variant deposition at promoters, underscoring a variant-specific role for SP1 in directing H2BE chromatin incorporation.

### The FACT complex restricts H2BE to the TSS

Mass spectrometry identified the histone chaperone FACT as interacting with H2BE. FACT is a well-characterized complex that facilitates H2A–H2B eviction and reassembly during RNA polymerase II elongation through nucleosomes^22,23,47^. In yeast and *Drosophila*, disruption of FACT complex activity leads to mis-localization of histone variants H2A.Z and CENP-A, suggesting an essential role for FACT in maintaining proper variant positioning during transcription^48–50^. We first validated the association between the FACT subunit SSRP1 and H2BE in both primary neurons and N2A cells (Figure S4A-B). Pharmacological inhibition of FACT using CBL0137^51^ (FACTi) did not alter total H2BE protein levels, indicating that FACT does not regulate H2BE expression or stability (Figure S4C).

To define FACT’s role in H2BE chromatin localization, we treated neurons with FACTi for 24 hours and performed CUT&Tag followed by sequencing. FACT inhibition resulted in a striking redistribution of H2BE away from TSSs and into gene bodies (Figure 4A). This redistribution was characterized by a significant increase in genic H2BE peaks accompanied by a concomitant loss of H2BE at TSSs (Figure 4B–F, S4D). Overall, neurons gained more than 1,500 genic H2BE peaks following FACT inhibition in genes related to synapse organization and signal transduction (Figure 4G, S4E). Stratification by gene expression revealed that the largest gains in genic H2BE occurred within highly and moderately expressed genes (Figure S4F-G), consistent with prior studies demonstrating that FACT promotes histone recycling and nucleosome reassembly during active RNA polymerase II elongation^52^. These findings suggest that FACT normally limits the improper deposition or spreading and accumulation of H2BE to genic regions, thereby maintaining its enrichment at TSSs.

**Figure 4.**
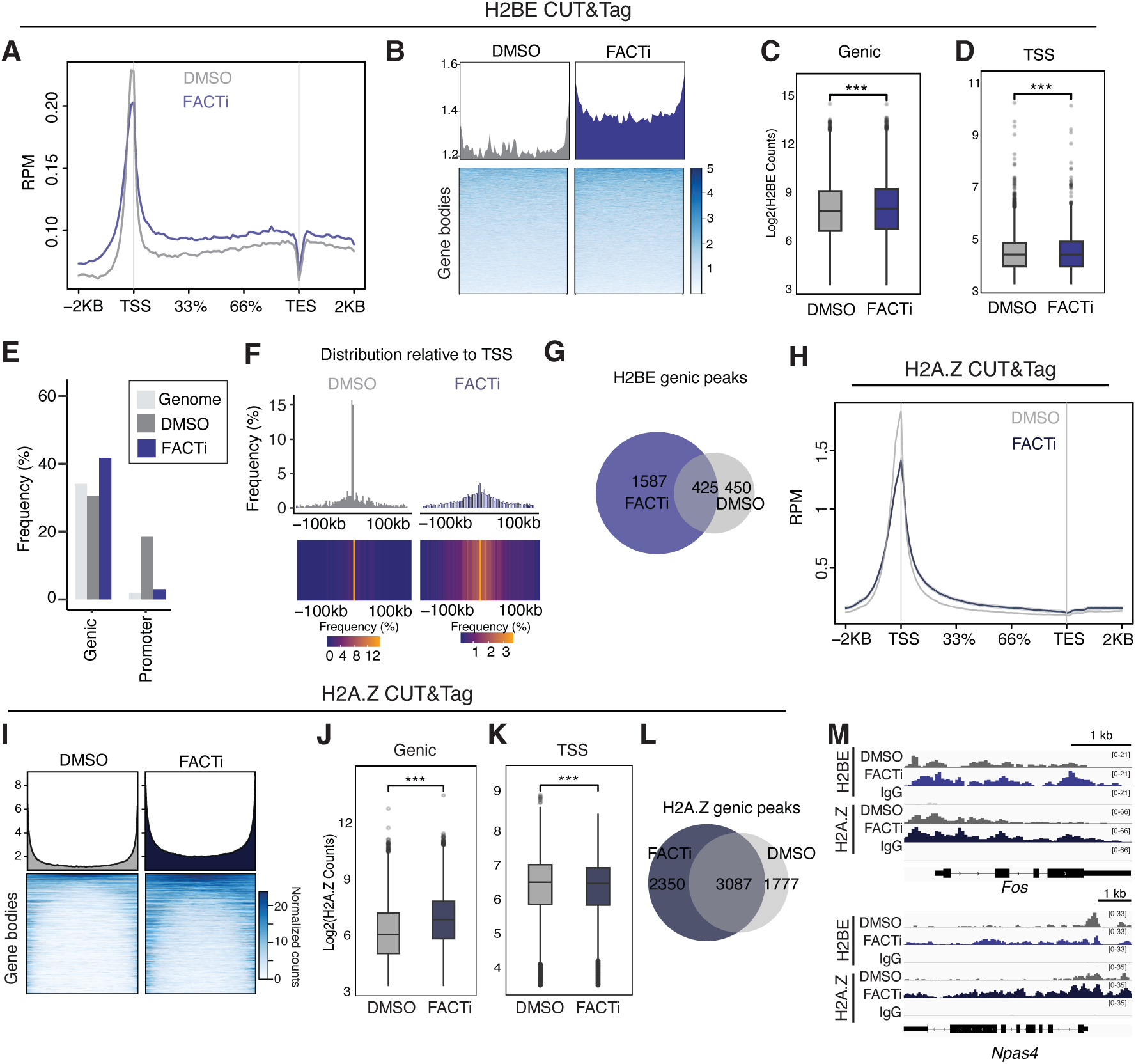
The FACT complex restricts H2BE to the TSS. A) Metaplot of H2BE CUT&Tag profiling in WT primary cortical neurons treated with DMSO (grey) or FACTi (blue). Plot shows read counts per million mapped reads (RPM) at ± 2 kb from the transcription start site (TSS) (*n* = 3 biological replicates per condition). B) Heatmap and metaplot depicting H2BE at gene bodies (genic regions minus first and last 500 bp). C) Normalized H2BE CUT&Tag read counts at H2BE peak sites with ≥ 10 reads detected at genic regions minus first and last 500 bp. D) Normalized H2BE CUT&Tag read counts at H2BE peak sites with ≥ 10 reads detected +/- 500bp from the TSS (unpaired t test, *** p <0.001). E) Genomic ranges analysis of the frequency of H2BE at genic or promoter regions. F) Genomic ranges analysis of the distribution of H2BE relative to the TSS in DMSO and FACTi conditions. G). Overlap of H2BE genic peaks called by MACS between DMSO and FACTi (peak score = 50). H) Metaplot of H2BE CUT&Tag profiling in WT primary cortical neurons treated with DMSO (grey) or FACTi (blue). Plot shows read counts per million mapped reads (RPM) at ± 2 kb from the transcription start site (TSS) (*n* = 3 biological replicates per condition). I) Heatmap and metaplot depicting H2A.Z at gene bodies (genic regions minus first and last 500 bp). J) Normalized H2A.Z CUT&Tag read counts at H2A.Z peak sites with ≥ 10 reads detected at genic regions minus first and last 500 bp. K) Normalized H2A.Z CUT&Tag read counts at H2A.Z peak sites with ≥ 10 reads detected +/- 500bp from the TSS (unpaired t test, *** p <0.001) Overlap of H2A.Z genic peaks called by MACS between DMSO and FACTi (peak score = 50). M) Representative gene tracks of genes that gained both an H2BE (top) and H2A.Z (bottom) genic peak with FACTi. Bigwigs were averaged across replicates.

To determine whether FACT’s role in H2BE localization is H2BE-specific in neurons, we performed CUT&Tag for H2A.Z under the same conditions. Consistent with observations in other models, FACTi similarly led to redistribution of H2A.Z away from TSSs and into gene bodies (Figure 4H–L). Notably, many genes that gained a genic H2BE peak upon FACT inhibition also gained a genic H2A.Z peak (Figure 4M), indicating that FACT acts broadly to constrain histone variant accumulation within gene bodies during transcription. Together, these results demonstrate that FACT functions as a transcription-coupled chaperone that restricts H2BE, among other histone variants, to promoter-proximal regions by preventing their aberrant incorporation or retention within gene bodies, thereby preserving proper variant localization during ongoing transcription.

### NAP1L4 evicts H2BE from chromatin

The dynamic regulation of chromatin, which is especially important in neurons for processes like synaptic plasticity and memory consolidation requires both histone variant incorporation and eviction^13,14,53^. We therefore examined H2BE-associated proteins (Figure 1D) to identify chaperones that may act to evict it from chromatin and identified NAP1L4 (NAP-1 like protein 4) as a shared binding partner between H2B and H2BE (Figure 1D-E). NAP-1 is a highly conserved histone chaperone involved in the dynamic regulation, including both incorporation^54^ and eviction^55,56^, of the H2A-H2B heterodimer. NAP1L4 has also been shown to associate with H2A variants, suggesting that it also plays a role in replication-independent chromatin maintenance^54^ but is less well understood than NAP-1.

To assess how NAP1L4 affects H2BE localization, we first validated the association between H2BE and NAP1L4 in N2A cells (Figure 5A). Next, we generated an shRNA lentivirus targeting *Nap1l4* and confirmed robust knockdown at both the mRNA and protein levels in N2As and primary cortical neurons (Figure S5A, B). Notably, loss of NAP1L4 resulted in an increase in total H2BE protein levels (Figure S5B), suggesting that NAP1L4 normally limits steady-state H2BE abundance, potentially by promoting histone eviction or turnover. Given this increase in total H2BE, we next asked whether NAP1L4 depletion alters the genomic distribution of H2BE. We transduced primary cortical neurons with a *Nap1l4* or nontargeting shRNA and used CUT&Tag to define its effects on H2BE localization. *Nap1l4* depletion significantly increased H2BE at the TSS compared to control (Figure 5B-D) and resulted in 2769 additional H2BE peaks identified above threshold (Figure 5E). Gained peaks were at the TSSs of genes related to metabolism, cell signaling, and cell adhesion (Figure S5C). Like other perturbations tested, we found that *Nap1l4* depletion affected genes of all expression levels, demonstrating a broad role for NAP1L4 in regulating H2BE chromatin association rather than an effect restricted to a specific transcriptional state (Figure S5D). Given that *Nap1l4* depletion leads to increased H2BE at the TSS, we compared these data to those using the transcriptional inhibitor Flavopiridol, which also caused increased H2BE at the TSS (Figure S3E-H). *Nap1l4* depletion caused a significantly greater increase in H2BE at the TSS than Flavopiridol treatment (Figure S5E), again suggesting that effects are not only due to disruptions to transcription. There was also little overlap between gained peaks (Figure S5F-G), indicating that these perturbations did not increase H2BE through similar mechanisms and that NAP1L4 action on H2BE is not likely due to an indirect effect on transcriptional disruptions.

**Figure 5:**
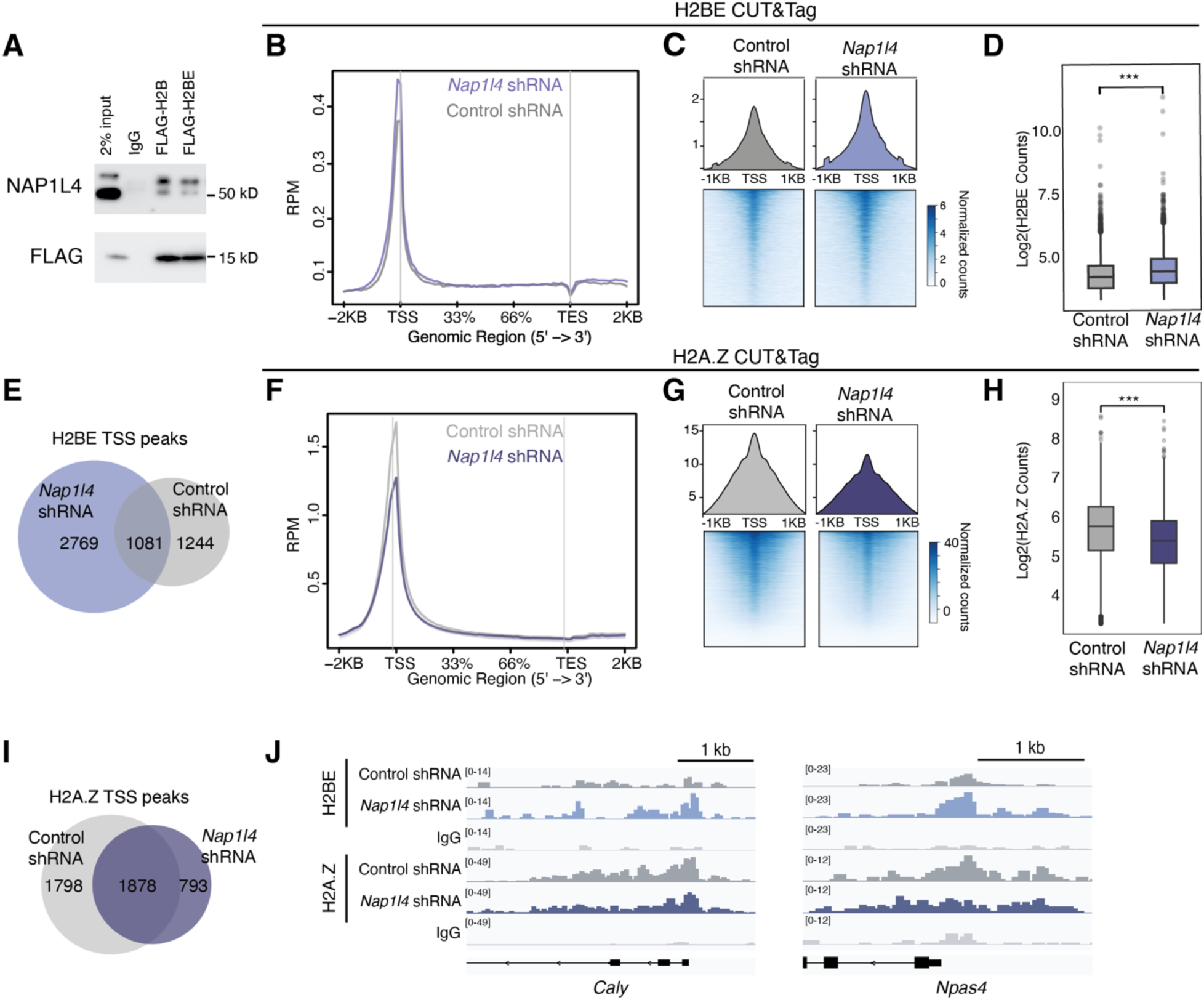
NAP1L4 evicts H2BE from chromatin. A) CoIP for FLAG-tagged H2B and H2BE and NAP1L4 in N2A cells. B) Metaplot of H2BE CUT&Tag profiling in WT primary cortical neurons transduced with a Control (grey) or *Nap1l4* shRNA (purple). Plot shows read counts per million mapped reads (RPM) at ± 2 kb from the transcription start site (TSS) (*n* = 2 biological replicates per condition). C) Metaplot and heatmap at +/- 1KB from the TSS of all genes for Control and *Nap1l4* depletion. D) Normalized H2BE CUT&Tag read counts at H2BE peak sites with ≥ 10 reads detected at the TSS (unpaired t-test). E) Overlap of H2BE TSS peaks called by SEACR between Control and *Nap1l4* shRNA. Peak score = 7. F) Metaplot of H2A.Z CUT&Tag profiling in WT primary cortical neurons transduced with a Control (grey) or *Nap1l4* shRNA (dark purple). Plot shows read counts per million mapped reads (RPM) at ± 2 kb from the transcription start site (TSS) (*n* = 3 biological replicates per condition). G) Metaplot and heatmap at +/- 1KB from the TSS of all genes for Control and *Nap1l4* depletion. H) Normalized H2A.Z CUT&Tag read counts at H2A.Z peak sites with ≥ 10 reads detected at the TSS (unpaired t-test). I) Overlap of H2A.Z TSS peaks called by SEACR between Control and *Nap1l4* shRNA. Peak score = 25. J) CUT&Tag gene tracks for representative genes. Bigwigs were averaged across replicates. H2BE is shown at the top, H2A.Z at the bottom

To determine whether NAP1L4’s function was specific to H2BE or reflected a broader role in histone variant eviction, we performed CUT&Tag for H2A.Z following *Nap1l4* depletion. In contrast to the increased accumulation of H2BE at TSSs, loss of NAP1L4 resulted in a significant decrease in H2A.Z enrichment at TSSs (Figure 5F–J). This opposing effect highlights a selective role for NAP1L4 in regulating histone variant occupancy, rather than acting as a general histone eviction factor. Taken together, these results suggest that NAP1L4 mediates the dynamic removal of H2BE at genes important for neuronal function and environmental response.

### Transcriptional consequences of H2BE chaperone depletion

Given that all components of the H2BE regulatory pathway characterized here, have with well-established roles in chromatin regulation beyond histone variant regulation, we next sought to define their H2BE-dependent transcriptional effects. Notably, even short-term inhibition of SP1 or FACT led to extreme transcriptional dysregulation, with each condition altering the expression more than 9,000 genes (Figure S6A-B). Such widespread changes likely reflect a dramatic shift in cellular state that could preclude interpretation of direct H2BE-dependent mechanisms. We therefore focused our transcriptional analyses on BAF and NAP1L4, whose opposing roles in H2BE incorporation and eviction offer a direct way to examine transcriptional consequences of bidirectionally altering H2BE dynamics.

#### BAF-mediated H2BE incorporation shapes neuronal gene expression programs

First, we asked how decreased incorporation of H2BE into chromatin resulting from BAF inhibition would affect gene expression. We performed RNA sequencing of WT and H2BE knockout (KO) primary cortical neurons that were treated with either DMSO or BAFi. In WT neurons, BAFi caused widespread transcriptional dysregulation, with most differentially expressed genes (DEGs) decreasing in expression (Figure 6A). Genes regulated by BAFi, particularly downregulated genes, were enriched for biological processes related to core neuronal identity terms, including regulation of synapse structure or activity, forebrain development, cognition, and learning (Figure S6C-D).

**Figure 6.**
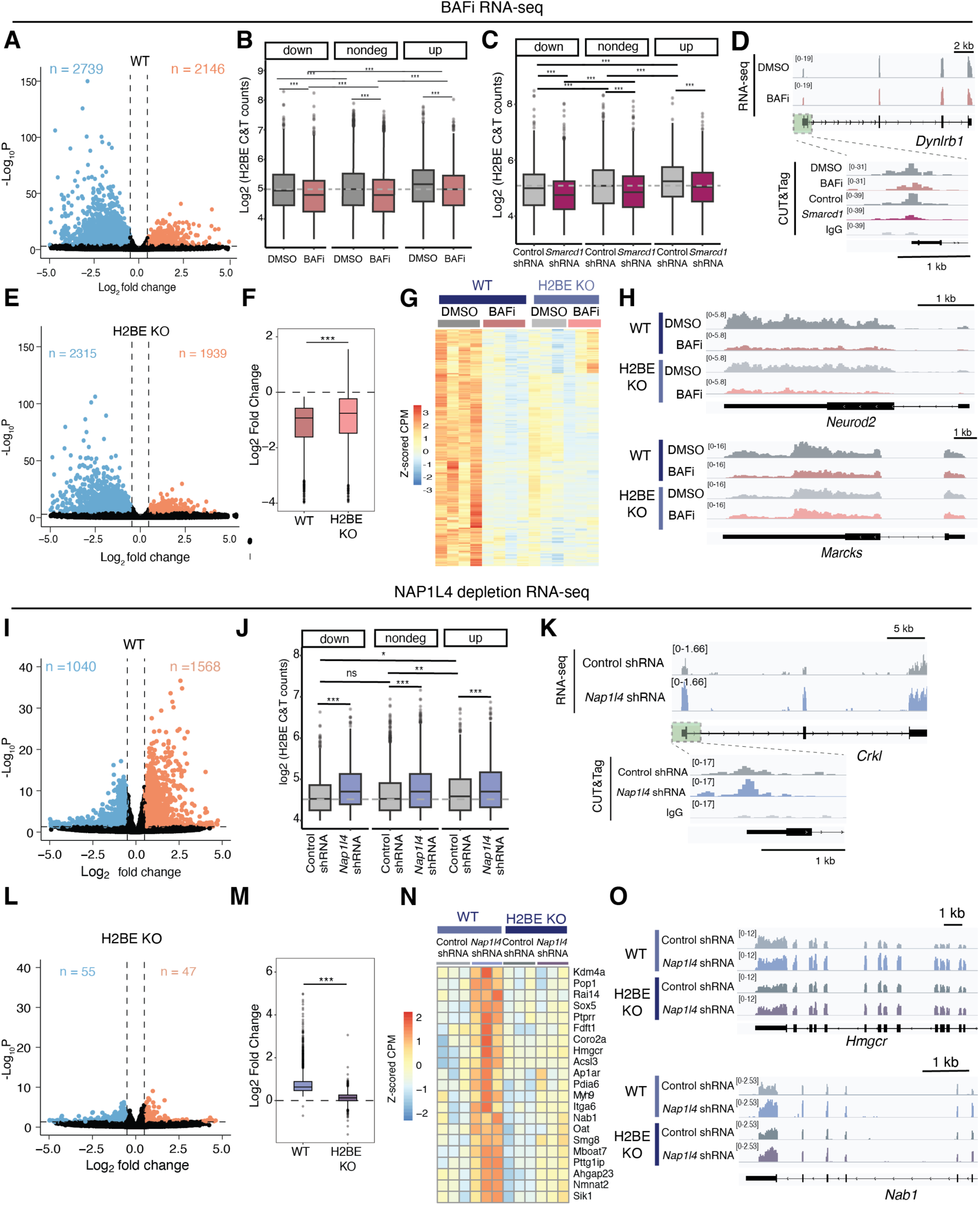
H2BE-dependent transcriptional effects of chaperone loss. A) Volcano plot of BAFi vs DMSO in WT neurons, n = 4 biological replicates/treatment. Vertical lines represent a log2-fold change cutoff of 0.5 (absolute value) and horizontal lines represent a p-value cutoff of 0.001. B) Log2 H2BE counts from BAFi H2BE CUT&Tag dataset binned by RNA-seq designation. Significance was calculated using a one-way ANOVA, followed by a post hoc Tukey’s HSD test. C) Same as B, this time using *Smarcd1* depletion H2BE CUT&Tag dataset. D) Representative gene track of a gene that both lost H2BE at its TSS with BAF perturbation and had decreased gene expression. For CUT&Tag, bigwigs were generated from merged replicates. For RNA-seq, tdfs were generated from individual replicates and then merged. E) Volcano plot of BAFi vs DMSO in H2BE KO neurons, n = 3 biological replicates/group. Vertical lines represent a log2-fold change cutoff of 0.5 (absolute value) and horizontal lines represent a p-value cutoff of 0.001. F) Boxplot of the log2fold changes (BAFi - DMSO) of genes downregulated by BAFi in WT neurons, shown in both H2BE KO and WT neurons. *** represents p < 0.001 in an unpaired t-test. G) Heatmap plotting the Z-scored counts per million of all genes that were downregulated by BAFi in the WT condition. H) Representative gene tracks of genes from the heatmap in F that were downregulated by both BAFi in WT neurons, and by H2BE KO. Tdfs were generated from individual replicates and then merged. I) Volcano plot showing differentially expressed genes in WT neurons comparing Control and *Nap1l4* shRNA. n = 3 biological replicates per genotype. Vertical lines represent a log2-fold change cutoff of 0.5 (absolute value) and horizontal lines represent a p-value cutoff of 0.001. J) Normalized H2BE CUT&Tag read counts at H2BE peak sites detected at the TSS for both Control and *Nap1l4* shRNAs, binned by whether the gene was down, unchanged, or upregulated in the RNA-seq shown in I. K) Representative gene track for a gene that had increased gene expression (top) and gained an H2BE peak (bottom) and with *Nap1l4* depletion compared to control. Bigwigs were merged across replicates. L) Volcano plot showing differentially expressed genes between H2BE primary KO cortical neurons transduced with control or *Nap1l4* shRNAs (*n* = 3 biological replicates per genotype). Vertical lines represent a log2-fold change cutoff of 0.5 (absolute value) and horizontal lines represent a p-value cutoff of 0.001. M) Boxplot of the log2fold changes (*Nap1l4* shRNA – Control shRNA) of genes upregulated by *Nap1l4* depletion in WT neurons, shown in both H2BE KO and WT neurons. *** represents p < 0.001 in an unpaired t-test. N) Heatmap of z-scored CPM of a subset of genes that are upregulated when *Nap1l4* is depleted in a WT context, but where that effect is dampened when *Nap1l4* is depleted down in an H2BE KO context. O) Tdfs were generated from individual replicates and then merged.

To define how H2BE deposition relates to transcriptional responses to BAFi, we integrated BAFi and *Smarcd1* depletion H2BE CUT&Tag datasets with the BAFi RNA-seq data in WT neurons. BAFi caused a global reduction of H2BE compared to control conditions across all gene expression groups in both CUT&Tag datasets (Figure 6B-C). Interestingly, we observed an inverse correlation between H2BE levels and the transcriptional effect of BAF inhibition. The genes with the lowest levels of H2BE upon BAFi inhibition, were downregulated upon BAFi. Conversely, genes that were upregulated retained the highest levels of H2BE following inhibition. This suggests that the dosage of H2BE may, in part, dictate sensitivity to BAFi such that genes with lower levels of H2BE subsequently lose enough following BAFi to be significantly downregulated. For example, genes such as *Dynlrb1*, a gene important for neurite outgrowth in sensory neurons^57^, both lost an H2BE peak and had decreased gene expression with BAFi (Figure 6D). However, some genes, such as *Atp1313* were upregulated by BAFi despite losing H2BE at the TSS (Figure S6E), likely due to a secondary effect of BAFi or other functions of the BAF complex in remodeling chromatin. Together, these findings demonstrate that some genes respond directly to changes in H2BE deposition caused by BAF inhibition, while others are governed by more complex regulatory mechanisms.

BAF complexes have established wide-ranging effects on chromatin beyond just the regulation of H2BE. To determine which subset of transcriptional effects of BAF inhibition depends on H2BE, we profiled gene expression in H2BE-KO neurons treated with DMSO or BAFi. As in WT neurons, downregulated DEGs (n = 2315) outnumbered upregulated DEGs (n = 1939) following BAF inhibition (Figure 6E), but the overall number of DEGs was lower in KO neurons. Globally, BAFi caused significantly less transcriptional downregulation in H2BE KO neurons compared to WT neurons (Figure 6F-G). This is consistent with a model in which BAF acts in part through promoting H2BE incorporation in neurons, and in which loss of H2BE attenuates some BAF-dependent transcriptional responses. Because H2BE promotes chromatin accessibility^17^, some of the dampened response in KO neurons may also be due to greater chromatin compaction preventing BAF complexes from acting. To more stringently test for genes in which BAF directly regulates expression through H2BE incorporation, we overlapped gene sets that were differentially by both BAFi and H2BE KO (Figure S6F, G). Based on a model that the BAF complex incorporates H2BE, we focused on the subset of 234 genes that were downregulated both by BAFi in WT neurons and in H2BE KO relative to WT (Figure S6F, H, 6G). For these genes, H2BE KO and BAFi decreased expression though not to the same degree, and the addition of BAFi in a KO context further decreased expression suggesting both shared and independent effects of BAFi (Figure 6G-H). These findings establish that BAF-regulated H2BE incorporation into chromatin contributes to neuronal transcriptional regulation.

#### H2BE-dependent consequences of *Nap1l4* depletion

Given the CUT&Tag data establishing NAP1L4’s role in H2BE eviction, we hypothesized that knocking down *Nap1l4* would increase gene expression. We conducted RNA-sequencing on WT and H2BE KO primary cortical neurons transduced with either a Control or *Nap1l4*-targeting shRNA. Consistent with this hypothesis, depletion of *Nap1l4* in WT neurons predominantly upregulated gene expression (Figure 6I), particularly of genes involved in synapses, axonogenesis, and dendrite development (Figure S6J). Downregulated genes were largely related to metabolic processes, including oxidative phosphorylation and mitochondrial translation (Figure S6K).

Integrating RNA-seq and H2BE CUT&Tag data revealed a direct correlation between H2BE occupancy at the TSS, and gene expression changes induced by *Nap1l4* depletion. *Nap1l4* depletion resulted in a global increase in H2BE across all differential gene expression groups, with genes upregulated by *Nap1l4* depletion showing the highest baseline levels of H2BE and a further upregulation following depletion (Figure 6J). This suggests that H2BE accumulation caused by *Nap1l4* depletion promotes transcriptional activation at these genes, consistent with H2BE’s role in promoting chromatin accessibility. For example, *Crkl*, a gene encoding an adapter protein important for transducing intracellular signals in the brain, gains H2BE at its TSS upon *Nap1l4* depletion, and is similarly upregulated in RNA-seq (Figure 6K).

Next, we investigated how *Nap1l4* depletion affects gene expression in H2BE KO neurons. We observed substantial blunting of transcriptional changes, with very few DEGs in KO neurons (55 down, 47 up, Figure 6L). Genes that were upregulated by *Nap1l4* depletion in a WT context were almost unchanged in the H2BE KO condition (Figure 6M) suggesting that some of NAP1L4’s roles are dependent on H2BE. As with BAF inhibition, because H2BE KO neurons have less open chromatin at baseline^17^, some of these effects may be due to NAP1L4 action directly on H2BE while others due to a more repressive chromatin state that could limit NAP1L4 function more broadly. To more thoroughly test for genes likely controlled by directly NAP1L4 regulation of H2BE, we focused on a subset of genes that showed a blunted effect of *Nap1l4* depletion in KO neurons but little baseline difference between WT and KO neurons (Figure S6L, 6N-O). For these genes, the absence of H2BE alone does not cause any overt changes in expression, suggesting that the lack of effects of *Nap1l4* depletion is not due to indirect effects of H2BE KO causing robust disruptions to chromatin at these loci. Rather, the divergent effects of *Nap1l4* loss at these loci in WT and KO neurons are more likely to be attributable to its role in H2BE eviction. Notably, many genes were regulated by *Nap1l4* depletion in WT neurons and similarly disrupted in H2BE KO neurons (Figure S6L-N), suggesting much of the blunted effect of *Nap1l4* depletion in KO neurons was due to baseline changes in gene expression. These results highlight that while NAP1L4 has some H2BE-dependent effects, it also exerts additional regulatory functions on chromatin. Together, these results define the transcriptional effects of the BAF complex and NAP1L4 in neurons and their role in with H2BE regulation, providing the first demonstration of their roles in controlling histone variant incorporation and eviction to modulate neuronal transcription.

## Discussion

Here, we identify binding partners for H2BE, including interactors shared with canonical H2B and those specific to H2BE. Building on these findings, we establish a model in which the BAF complex, guided by the transcription factor SP1, deposits H2BE, while NAP1L4 removes it. These opposing regulatory mechanisms shape H2BE-dependent gene expression. In addition, the FACT complex maintains H2BE at TSSs of genes by counteracting its spread and subsequent accumulation in genic regions. Comparison with the H2A variant H2A.Z reveals that these mechanisms include both shared and H2BE-specific pathways, highlighting how chromatin machinery can fine-tune nucleosome composition. Together, these findings define the first regulatory network for the localization of H2BE in neurons and demonstrate how multiple chaperone proteins and transcription factors can work in concert to regulate variants that only subtly differ from canonical histones (Figure 7).

**Figure 7.**
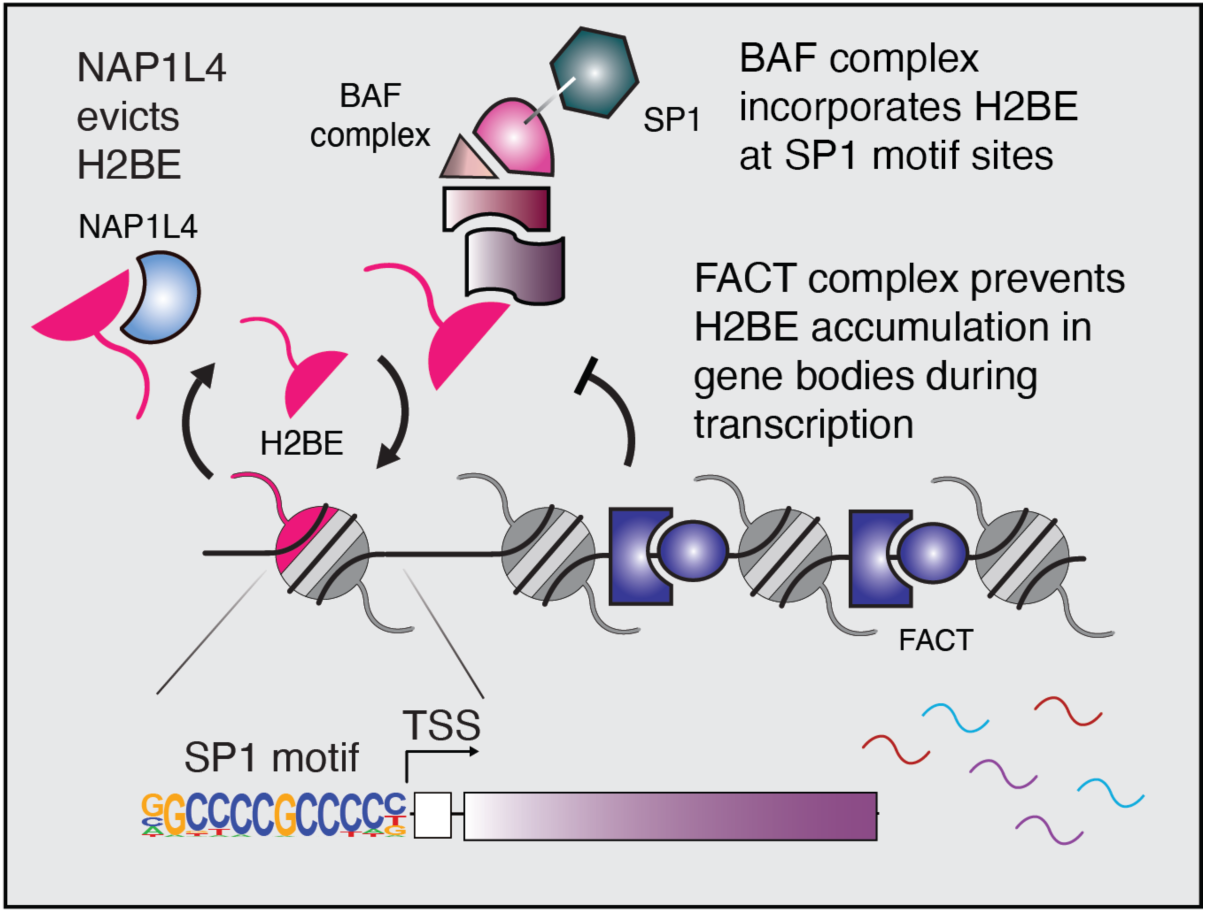
Model for H2BE regulation in neuronal chromatin. Guided by the transcription factor SP1, the BAF complex incorporates H2BE at the TSS of genes with SP1 motif sites. The FACT complex restricts H2BE from accumulating in gene bodies during transcription. NAP1L4 evicts H2BE from chromatin, promoting variant exchange. Together, these factors dictate H2BE localization in the genome and effects on transcription.

Biochemical analysis of H2BE binding partners via MS revealed unexpected differences in H2B and H2BE-containing nucleosomes. Notably, the lack of association between H2BE and canonical H2B in these data suggests that H2BE-containing nucleosomes are likely homotypic. Given the extremely low abundance of H2BE compared to canonical H2B (<1%), this suggests a remarkable degree of precision in directing H2BE to the correct nucleosomes, since it would be unlikely for cells to achieve H2BE homotypic nucleosomes by chance. Also noteworthy, we found decreased association between H2BE and H1 linker histones and MacroH2A compared to canonical H2B. The diminished association between H2BE and MacroH2A is notable given that MacroH2A stabilizes the nucleosome and impedes BAF complex recruitment and remodeling^26,58^. Consistent with a model in which H2BE promotes chromatin accessibility and is deposited by the BAF complex, the repressive architecture imposed by MacroH2A could act as a barrier to H2BE incorporation.

Despite these differences, most identified chaperones were shared between H2B and H2BE. This observation raises questions about whether established models of chaperone variant function are relevant to H2B variants. It has long been hypothesized that every histone variant possesses a unique, dedicated chaperone^59^. However, other histone variants like H3.3, MacroH2A, and H2A.Z contain motifs distinct from their canonical counterparts, facilitating recognition by their specific chaperones. While H2BE does differ from H2B, it does not contain specific motifs, but rather 5 unique amino acids spread throughout the protein^16^. Many other distinct H2B encoding genes also only differ very slightly from canonical H2B, often by only a single amino acid^60^. The highly similar sequences between canonical H2B and H2BE, as well as other putative H2B variants, suggest that the H2B family may offer a unique case in which chaperone proteins are not specific to variants but rather shared with the broader H2B family.

Consistent with this idea, our data support a model in which H2BE achieves specificity not through exclusive chaperones, but through the activity of multiple complexes paired with 1) specific targeting by transcription factors to active regions, 2) the ongoing generation of H2BE in non-dividing cells making it available for incorporation at sites of high accessibility and nucleosome turnover, and 3) its ability to open chromatin which may allow for more selective loss from specific genomic regions. Such a model could position H2BE at sites of active transcription where transcription factors are targeting and chromatin remains highly accessible. We propose that while BAF may bind multiple H2B species, most are ‘canonical’ and thus not continually generated in post-mitotic cells. Thus, the continued generation of H2BE paired with cooperation with transcription factors like SP1 might mediate the specific incorporation of H2BE at active sites that undergo regular nucleosome turnover and require continued histone replacement and greater accessibility. Similarly, exchange with canonical H2B by shared binding partners such as NAP1L4 and FACT may be supported by H2BE’s ability to open chromatin, causing a less stable chromatin complex that pushes the reaction towards eviction. Lastly, some degree of specific deposition may simply be achieved by histone exchange being most likely to occur in regions of open chromatin and active transcription, allowing H2BE to support a positive feedback loop that promotes highly accessible chromatin.

A central finding of this work is that the BAF complex promotes the incorporation of H2BE into chromatin. While BAF is primarily recognized as an ATP-dependent remodeler, our data support an emerging role for this complex as a histone chaperone. Subunits within ATP-dependent chromatin remodeling complexes often possess histone-binding domains that facilitate histone exchange during the remodeling cycle^61,62^. Consistent with this, BAF has been shown to facilitate H2A-H2B dimer exchange^63^, a mechanism mirrored by other remodelers like SWR1/SCRAP, which deposits H2A.Z^8,30,64^. We propose a model in which BAF-mediated chromatin remodeling generates a transiently accessible nucleosome state that allows displacement of canonical H2A-H2B dimers and replacement with H2A-H2BE dimers when available. Importantly, this activity is spatially constrained through cooperation with the transcription factor SP1. SP1 motifs are highly enriched at promoters that accumulate H2BE, and SP1 physically associates with both BAF and H2BE. Loss of SP1 leads to aberrant, genome-wide expansion of H2BE deposition in a BAF-dependent manner, indicating that SP1 ordinarily restricts BAF-mediated H2BE incorporation to selected promoters. Transcription factor mediated recruitment of the BAF complex has been described previously^65^, establishing a framework in which sequence-specific factors target chromatin remodeling activity. Analogous recruitment mechanisms have been reported for H3.3 and H2A.Z^66,67^, highlighting how transcription factors can specify histone variant deposition to modulate chromatin. Notably, *Sp1* depletion did not induce comparable expansion of H2A.Z at promoters, underscoring the specificity of this mechanism for H2BE. Thus, despite acting on shared chromatin substrates, BAF and SP1 together can generate variant-specific outcomes, highlighting how transcription factor context can sculpt histone variant landscapes.

We also found that multiple proteins contribute to the removal of H2BE from chromatin. Depletion of *Nap1l4* caused a pronounced accumulation of H2BE at TSSs, exceeding that observed upon acute transcriptional inhibition and affecting largely distinct gene sets. This indicates that NAP1L4 does not regulate H2BE indirectly through transcriptional repression but instead promotes its eviction from chromatin. Strikingly, *Nap1l4* depletion produced the opposite effect on H2A.Z, leading to decreased H2A.Z occupancy at promoters. This divergent response demonstrates that NAP1L4 is not a general H2A–H2B eviction factor, but instead differentially regulates histone variant composition. While NAP-1 family proteins are classically associated with H2A-H2B shuttling^54–56^, our findings indicate that NAP1L4 has a specialized role in controlling H2BE dynamics in neurons. Given that H2BE removal is essential for neuronal homeostatic plasticity^19^, NAP1L4 is also well positioned to function as a key regulator of experience-dependent chromatin remodeling. However, whether NAP1L4 directly exchanges H2BE for canonical H2B or promotes eviction through indirect destabilization of H2BE-containing nucleosomes remains an important question for future studies.

Our data further identify FACT as a critical regulator of H2BE localization. Inhibition of FACT resulted in redistribution of H2BE away from TSSs and into gene bodies, suggesting FACT is required to prevent the accumulation of H2BE in genic regions. To determine whether this function was specific to H2BE, we examined the localization of H2A.Z following FACT inhibition. Like H2BE, FACT inhibition caused mis-localization of H2A.Z into gene bodies, a phenotype previously reported in yeast FACT mutants^48^. As is the case with H2A.Z^48^, loss of FACT might cause improper deposition of H2BE by its other chaperones like the BAF complex. Alternatively, FACT might actively remove H2BE-containing nucleosomes that have spread into the gene body. Together, these data position FACT as a critical regulator of both H2BE and H2A.Z, reinforcing its broader role in maintaining chromatin architecture during transcription.

### Limitations of the study

While the findings presented here define multiple novel regulatory mechanisms that control a critical histone variant protein, much remains to be discovered. The effect sizes observed in CUT&Tag may be an underestimate of the true regulatory impact of BAF, SP1, FACT and NAP1L4 due to partial depletion of these proteins. Alternatively, these effects could suggest functional redundancy. For example, multiple chaperones regulate H2A.Z, underscoring the biological importance of ensuring variant availability^62,68^. Whether H2BE utilizes a similarly redundant network of chaperones remains to be determined. Similarly, the mechanism by which FACT restricts H2BE from the TSS remains to be explored, as it may act by preventing the deposition by other chaperone proteins such as BAF, or by directly removing H2BE that spreads into genic regions as it processes through the gene body. Further, mass spectrometry identified specific interactions between H2BE and other proteins of interest, only a subset of which we were able to explore. Notably, we found that H2BE interacts with other potential chaperones such as ANP32E, NPM1, and NCL complex proteins.

ANP32E regulates steady-state H2A.Z occupancy in neurons, with effects on transcriptional output, dendritic morphology, and memory formation^53^. ANP32E may contribute more broadly to H2B variant regulation in neurons, either directly or by shaping chromatin environments permissive for variant exchange. These findings further support a model in which H2B variant localization is governed by shared chaperone pathways rather than strictly variant-specific factors. Finally, mass spectrometry also detected several proteins important for RNA processing and metabolism. While the functional significance of these interactions remains unknown, these and other unexplored binding partners identified here represent a rich dataset for future investigation into the transport, RNA-association, and post-translational modification of H2BE. In sum, we present the first evidence for the regulation of H2BE in chromatin, identifying that the BAF complex incorporates H2BE at TSSs under the guidance of SP1, FACT maintains H2BE at TSSs, and NAP1L4 mediates H2BE eviction from chromatin. This dynamic positioning and exchange of H2BE by chaperones offers a new framework for understanding how it modulates chromatin dynamics and transcription.

## Resource availability

### Lead contact

Requests for further information, resources, and reagents should be directed to and will be fulfilled by the lead contact, Erica Korb.

### Materials availability

This study did not generate new, unique reagents.

### Data and code availability

This paper analyzes existing, publicly available data, available at: GSE245256.

## Acknowledgements

This work was supported by the National Institutes of Health grant 1DP2MH129985 (EK), National Institutes of Health grant R01NS134755 (EK), and the Blavatnik Family Fellowship in Biomedical Research (AKS). We would like to thank the Children’s Hospital of Philadelphia Proteomics Core for services performed and assistance with analysis. Thanks also to Drs. Kenneth Zaret and Zhaolan Zhou for their critical reading of the manuscript.

## Author Contributions

Conceptualization: EK; investigation: AKS, ERF, RPT; visualization: AKS; formal Analysis: AKS; supervision: EK; writing – original draft: AKS; writing – review and editing: EK; funding acquisition: EK.

## Declaration of generative AI and AI-assisted technologies in the writing process

During the preparation of this work the authors used Microsoft Copilot to improve grammer and clarity of the manuscript after an initial draft was written. After using this tool, the authors reviewed and edited the content as needed and take full responsibility for the content of the published article.

## Methods

### Antibodies

Rabbit anti-H2BE: Millipore ABE1384

Rabbit anti-H2B: abcam ab1790

Rabbit anti-SMARCD1: Proteintech 10998-2-AP

Rabbit anti-SMARCC2: Cell Signaling Technology #8829

Rabbit anti-SMARCB1: Cell Signaling Technology #8745

Rabbit anti-NAP1L4: Proteintech 27889-1-AP

Rabbit anti-SP1: Millipore 07-645

Rabbit anti-H2A.Z: abcam ab4174

Goat Anti-Rabbit IgG, HRP: abcam ab6721

Guinea Pig anti-Rabbit IgG: Antibodies-online ABIN101961

### Mice

An H2BE-KO mouse was generated as described previously^16^. Briefly, the endogenous H2BE coding sequence was replaced with a membrane-targeted mCherry reporter sequence in C57Bl/6 mice. All mice were housed in a 12-hour light-dark cycle and fed a standard diet. Experiments were conducted in accordance with and approval of the IACUC.

### Cell Culture

#### Primary neuronal culture

Cortices were dissected from E16.5 C57BL/6J embryos and cultured in supplemented neurobasal medium (Neurobasal [Gibco 21103-049], B27 [Gibco 17504044], GlutaMAX [Gibco 35050- 061], Pen-Strep [Gibco 15140-122]) in TC-treated 12, 6-well or 10cm plates coated with 0.05 mg/mL Poly-D-lysine (Sigma-Aldrich A-003-E). At day in vitro (DIV) 3, neurons were treated with 0.5 μM AraC. Neurons were collected at DIV 12 for all experiments.

#### Neuro-2A (N2A) cells

Neuro-2A cells were obtained from the American Type Culture Collection (ATCC), cultured in DMEM (with 4.5 g L–1 glucose, L-glutamine and sodium pyruvate) supplemented with 10% FBS (Sigma-Aldrich, F2442-500ML) and 1% penicillin-streptomycin (Gibco, 15140122) and maintained free of mycoplasma.

#### Chaperone depletion/inhibition

##### shRNAs

*Smarcd1, Sp1* and *Nap1l4-tageting* and control Luciferase shRNAs were inserted into the pLKO.1 vector backbone (Addgene plasmid #10878). shRNA target sequences were as follows:

1. *Smarcd1* shRNA: GCCTGAAATCAAACGGGTAAA
2. *Sp1* shRNA 2: CCAATGCCAATAGTTATTCAA
3. *Nap1l4* shRNA: GCAGCTTTGCAGGAACGTCTT

##### Inhibition

BAFi (*BRD-K98645985* [MedChemExpress HY-114268]) was reconstituted in DMSO. Neurons were treated with 10μM. FACTi (CBL0137 [MedChemExpress HY-18935]) was reconstituted in DMSO. Neurons were treated with 2μM. Flavopiridol hydrochloride hydrate (Milipore F3055) was reconstituted in DMSO. Neurons were treated with 3μM. SP1i (Mithramycin A [MedChemExpress HY-A0122]) was reconstituted in DMSO and neurons were treated with 400nM.

#### Lentiviral production

HEK293T cells were cultured in high-glucose DMEM growth medium (Corning 10-013-CV), 10% FBS (Sigma-Aldrich F2442-500ML), and 1% Pen-Strep (Gibco 15140-122). Calcium phosphate transfection was performed with Pax2 and VSVG packaging plasmids. Viral media was removed 16 h after transfection and collected at 48 and 72 h later. Viral media was passed through a 0.45-μm filter and precipitated for 48 hours with PEG-it solution (40% PEG-8000 [Sigma-Aldrich P2139-1KG], 1.2 M NaCl [Fisher Chemical S271-1]). Viral particles were pelleted and resuspended in 200μL PBS.

#### Neuronal transduction

At DIV 5, neurons were transduced overnight with lentivirus containing the shRNA constructs described above. Virus was removed the following day, and neurons were cultured for six additional days.

##### Coimmunoprecipitations

Primary cortical neurons (1.2E6 cells/10cm dish) or N2A cells (8.8E6 cells/10cm dish) were washed with ice-cold sterile phosphate-buffered saline (PBS) and harvested in 1mL of Lysis Buffer 1 (50mM HEPES-KOH, pH 7.5-8, 140 mM NaCl, 1mM EDTA, 10% glycerol, 0.5% NP-40, 0.25%, Triton X-100, in ddH2O supplemented with 1X protease and phosphatase inhibitors, 1mM DTT and 1mM PMSF). The cell suspension was rotated for 10min at 4C and then centrifuged at 1350 x g for 5 mins at 4C. The pellet was resuspended in 1mL of LB2 (10mM Tris-HCl pH 8.0, 200mM NaCl, 1mM EDTA, 0.5mM EGTA in ddH2O) with inhibitors and rotated for 10 minutes at room temperature, centrifuged for 5 minutes at 1350 x g, then resuspended in LB3 (10mM Tris-HCl pH 8.0, 100mM NaCl, 1mM EDTA, 0.5mM EGTA, 0.1% Na-Deoxycholate, 0.5% N-lauroylsarcosine in ddH2O) with inhibitors. Note for CoIPs using a FLAG-tagged construct, LB3 was modified to 10mM Trish PH 8.0, 1mM EDTA, 0.5mM EGTA, 0.1% sodium-deoxyholate, 0.5% sodium lauroyl sarconsinate, 1% TritonX-100, 10mM sodium butyrate, 10mM nicotinamide. Chromatin and protein complexes were solubilized via sonication using a tip sonicator. Samples were maintained on ice throughout. Triton X-100 was added to a final concentration of 1% and the lysate was cleared by centrifugation at maximum speed for 10 min at 4C. 50uL (5%) input was collected from the supernatant and stored at -80C. The remaining supernatant (950uL) was reserved for the IP.

Magnetic Dynabeads Protein A (Invitrogen 10002D) were prepared by washing three times in block solution (0.5% BSA in PBS). For each IP, 50uL of beads were incubated with 1ug/mL of antibody in 500uL of block solution and coupled at 4C for a minimum of 2 hours. Antibody-bound beads were washed 3x in block solution on a magnetic rack and resuspended in 55uL of block solution. The prepared beads were added to 950uL of cleared lysate and incubated overnight at 4C with rotation. Following incubation, the bead-protein complexes were washed 3x with LBS and supplemented with 0.05% NP-40 and once with LB3 alone to remove non-specific binding.

### Quantitative Mass spectrometry

#### In-Solution Digestion

Immunoprecipitated proteins were eluted from the beads by resuspension in 0.3% SDS followed by water bath sonication for 5 minutes. Beads were separated on a magnetic rack and eluate was transferred to a new tube for acetone precipitation. Eluates were precipitated by the addition of 5M NaCl to a final concentration of 0.1M and 4 volumes of acetone. After mixing well, samples were incubated at room temperature for 30 minutes. Samples were centrifuged at 18000g for 10 minutes, after which the supernatant was removed. Protein pellets were then digested using the iST kit (PreOmics GmbH, Martinsried, Germany) per the manufacturer’s protocol^69^. Briefly, proteins were solubilized, reduced, and alkylated by the addition of SDC buffer containing TCEP and 2-chloroacetamide followed by heating to 95C for 10 minutes.

After cooling, samples were transferred to iST cartridges and digestion buffer containing LysC and trypsin was added. Samples were digested at 37C for 1.5 hours, after which the resulting peptides were washed and eluted from the cartridges. Eluates were dried by vacuum centrifugation and reconstituted in 0.1% TFA containing iRT peptides (Biognosys, Schlieren, Switzerland) for subsequent LC-MS analysis.

#### Mass Spectrometry Data Acquisition

Samples were analyzed on a Q-Exactive HF mass spectrometer (Thermofisher Scientific, San Jose, CA) coupled with an Ultimate 3000 nano UPLC system and an EasySpray source. Peptides were loaded onto an Acclaim PepMap 100 75um x 2cm trap column (Thermo) at 5uL/min and separated by reverse phase (RP)-HPLC on a nanocapillary column, 75 μm id × 50cm 2um PepMap RSLC C18 column (Thermo). Mobile phase A consisted of 0.1% formic acid and mobile phase B of 0.1% formic acid/acetonitrile. Peptides were eluted into the mass spectrometer at 300 nL/min with each RP-LC run comprising a 90-minute gradient from 3% B to 45% B.

Data was collected in Data Dependent Acquisition (DDA) mode with settings as follows: one full MS scan at a resolution of 120,000 ranging from 300-1800m/z, with an AGC target of 500,000 and a maximum injection time of 120ms. This was followed by data dependent MS2 scans on the twenty most abundant ions at a resolution of 45,000 and scan range from 200-2000m/z with an AGC target of 100,000, a maximum injection time of 120ms, and a collision energy of 27. Dynamic exclusion was set with a repeat count of 1 and duration of 15s. Unassigned, 1, 6-8, and >8 charge states were rejected.

#### System Suitability and Quality Control

As a measure for quality control, a standard *E. coli* protein digest was injected prior to and after the sample set and DDA data was collected. Resulting data were monitored using QuiC software (Biognosys, Schlieren, Switzerland) and searched in MaxQuant (Max Planck Institute, Martinsreid, Germany)^70^. MaxQuant output was visualized using the PTXQC^71^ package to track system performance and suitability.

#### Mass Spectrometry Raw Data Processing

DDA raw files were searched with MaxQuant (v.1.6.0)^70^ against the UniProt mouse reference database (UP000000589) containing reviewed sequences and isoforms and appended with sequences for common protein contaminants. The enzyme was specified as trypsin with a maximum of 2 missed cleavages. Carbamidomethylation of cysteine was specified as a fixed modification, and protein N-terminal acetylation and oxidation of methionine were considered variable modifications. A false discovery rate limit of 1% was set for peptide and protein identification. Remaining search parameters were kept as default values. Searched data were loaded into Scaffold (v.5.3.4) for visualization.

##### Subcellular fractionations

Protocol was adapted from^72^. Briefly, 8E6 N2A cells were washed twice in ice cold DBPS. Cells were collected in 1mL 25 mM HEPES pH 7.6, 25 mM KCl, 5 mM MgCl_2_, 0.05 mM EDTA, 0.1% NP-40, and 10% glycerol + protease inhibitors and rotated at 4C for 10 minutes. Nuclei were isolated by centrifugation at 6000 x g for 5 mins at 4C. 200uL of supernatant (cytoplasmic fraction) were transferred to a fresh tube. Pellet was then resuspended in 200uL of 100 mM Tris pH 8.0, 2% NP-40, 0.5% sodium deoxycholate, and 100mM of NaCl + protease inhibitors, resuspended, rotated at 4C for 30 mins, centrifuged for 3 mins at 6500 x g at 4C. 200uL of supernatant (nuclear fraction) was transferred to a fresh tube. The pellet was then resuspended in 200uL of 100 mM Tris pH 8.0, 2% NP-40, 0.5% sodium deoxycholate, and 1M of NaCl + protease inhibitors, and collected (chromatin fraction). All samples were sonicated for at least 5 minutes in a bioruptor (Diagenode). The nuclear fractions were sonicated for an additional 5 minutes (10 minutes total), and chromatin an additional 10 minutes (15 minutes total).

##### Western blotting

Protein lysates were mixed with 5X Loading Buffer (5% SDS, 0.3M Tris pH 6.8, 1.1mM Bromophenol blue, 37.5% glycerol), boiled for 10 minutes, and cooled on ice. Lysates were sonicated for 5 minutes in a bioruptor (Diagenode). Protein was resolved by 4 - 20% Tris-glycine SDS-PAGE, followed by transfer to a 0.45-μm PVDF membrane (Sigma-Aldrich IPVH00010) for immunoblotting. Membranes were blocked for 1 hour at RT in 5% milk in 0.1% TBST and probed with primary antibody overnight at 4C. Membranes were incubated with secondary antibody for 1 hour at RT.

##### CUT&Tag-sequencing

###### Library preparation & sequencing

Input samples were 500K primary cortical neurons per biological replicate. CUT&Tag was performed according to published protocols^34^. Concanavalin A-coated beads (EpiCypher 21-1411) were washed twice with 1mL cold filter-sterilized Binding Buffer (20mM HEPES pH 7.9, 10mM KCl, 1mM CaCl_2_, 1mM MnCl_2_) and resuspended in 11uL/reaction Binding Buffer. Cells were collected in 500uL room temperature Wash Buffer (20mM HEPES pH 7.5, 150mM NaCl, 0.5mM spermidine, supplemented by EDTA-free protease inhibitor [Roche 4693159001]), pelleted at 600 g for 3 min at room temp., washed once with 500uL Wash Buffer (room temp.), pelleted again, and resuspended in 100uL/reaction Wash Buffer (room temp.). To bind cells to ConA beads, 10uL activated beads were added to 100uL cells per reaction, vortexed briefly, and incubated for 10 min at room temperature. Beads were collected on a magnet, supernatant discarded, and resuspended in 50uL ice-cold Antibody Buffer (0.05% digitonin, 2mM EDTA, 0.1% BSA in Wash Buffer). To each reaction, 1ug primary antibody against the target protein was added (anti-H2BE [Millipore ABE1384], anti-H2A.Z [Abcam ab4174,] or rabbit IgG [Sino Biological CR1], vortexed briefly, and incubated overnight at 4°C on a rocking shaker. Beads were collected on a magnet, supernatant discarded, and resuspended in 50uL secondary antibody (guinea pig anti-rabbit IgG (antibodies-online ABIN101961, diluted 1:100 in Dig-Wash Buffer [0.05% digitonin in Wash Buffer]). Samples were then incubated for 1hr at room temperature on a nutator. On a magnet, beads were washed 2X with 200uL Dig-Wash Buffer. Beads were collected on a magnet, supernatant discarded, and beads were resuspended in 50uL loaded pA-Tn5 adapter complex (Diagenode C01070001-T30, diluted 1:250 in Dig-300 Buffer [20mM HEPES pH 7.5, 300mM NaCl, 0.5mM spermidine, 0.01% digitonin]) or 50uL loaded pG-Tn5 adapter complex^73^ and diluted 1:100 in Dig-300 buffer. Samples were then incubated for 1hr at room temperature on a nutator. On a magnet, beads were washed 2X with 200uL Dig-300 Buffer. Beads were collected on a magnet, supernatant discarded, and beads were resuspended in 300uL Tagmentation Buffer (10mM MgCl_2_ in Dig-300 buffer). Samples were then incubated for 1hr in a 37°C heat block. To stop tagmentation, 10uL 0.5M EDTA, 3uL 10% SDS, and 2.5uL 20mg/mL Proteinase K were added to each reaction and mixed by vortexing. Protein was digested for 1hr in a 55°C heat block. DNA was isolated by the DNA Clean & Concentraor-5 (Zymo Research D4014). A universal i5 primer and uniquely barcoded i7 primer were ligated and libraries were amplified by PCR with NEBNext High-Fidelity 2× PCR Master Mix (NEB M0541). Library clean-up was performed with AMPure XP beads (Beckman A63880) and eluted off beads in 25uL Tris-HCl pH 8. Prior to sequencing, library size distribution was confirmed by capillary electrophoresis using an Agilent 4200 TapeStation with high sensitivity D1000 reagents (5067-5585), and libraries were quantified by qPCR using a KAPA Library Quantification Kit (Roche 07960140001). Libraries were sequenced on an Illumina NextSeq1000 instrument (61-bp read length, paired end).

###### Data processing and analysis

Reads were mapped to *Mus musculus* genome build mm10 with Bowtie 2 (v2.5.1)^74^. Each biological replicate was subset to the same number of reads, and each condition was then merged across biological replicates (SAMtools v1.16.1)^75^. Heatmaps were generated using deepTools^76^d (v3.5.1) with a bedfile of the 500bps +/- the transcription start site. An additional 500bp were plotted on either end of the TSS. Metaplots were generated using ngs.plot^77^ (v2.63) against the mouse genome. For Figure 2 (BAFi and *Smarcd1* depletion), high confidence H2BE peaks were defined as sites that had ≥ 60 H2BE reads in the control condition. Peaks were called using SEACR^78^, stringent mode, normalized to IgG and annotated using Homer^79^ (v5.1). For the BRG1 published dataset^39^, MACS^80^ (v2.2.7.1) was used to call peaks, and a peak cutoff of 13 was applied.

IGV^81^ (v2.4.6) was used to generate genome browser views. Bigwigs were merged across replicates. To compare CUT&Tag signal to gene expression, normalized read counts from RNA-sequencing of WT primary neuronal cultures from^17^ were used to generate gene lists by expression level. Genes with base mean < 3 were defined as ‘not expressed’. The remaining genes comprise the ‘all expressed’ categorization, and these genes were further divided into 3-quantiles (by base mean) to define ‘low expression’, ‘mid expression’ and ‘high expression’.

###### Gene ontology

For gene ontology analysis from CUT&Tag, gene names were assigned to peak coordinates using Homer. PANTHER^82^ (v18.0) was used to perform an overrepresentation test against the biological process complete ontology using default parameters. Genes expressed in mouse neurons^17^ were used as the background list.

##### RNA-sequencing

###### Library preparation & sequencing

For BAFi RNA-seq, input samples were 200K primary cortical neurons from 4 WT biological replicates and 3 KO biological replicates. For *Nap1l4* depletion, input samples were 300K primary cortical neurons from 3 WT biological replicates and 3 KO biological replicates. For all, RNA was isolated using Quick-RNA Miniprep Plus Kit (Zymo Research R1057). Prior to library preparation, RNA integrity was confirmed using an Agilent 4200 TapeStation with high sensitivity RNA reagents (5067-5579). Sequencing libraries were prepared using the NEBNext Ultra II RNA Library Prep Kit (E7770). Prior to sequencing, library size distribution was confirmed by capillary electrophoresis using an Agilent 4200 TapeStation with high sensitivity D1000 reagents (5067-5585), and libraries were quantified by qPCR using a KAPA Library Quantification Kit (Roche 07960140001). Libraries were sequenced on an Illumina NextSeq1000 instrument (61-bp read length, paired end).

###### Data processing and analysis

Reads were mapped to *Mus musculus* genome build mm10 with Star^83^(v2.7.9a). The R package DESeq2^84^ (v1.42.1) was used to perform differential gene expression analysis. We defined genes as differentially expressed where adjusted p-value ≤ 0.01 and absolute fold change ≥ 0.5. Volcano plots were generated using EnhancedVolcano^85^ (1.26.0). IGV^81^ (v2.4.6) was used to generate genome browser views.

###### Gene ontology

For gene ontology analysis from RNA-sequencing, clusterProfileR^86^ (v4.18.2) was used. Genes expressed in mouse neurons were used as the background list.

## Figure Legends

**Supplementary Figure 1. Identification of H2BE binding partners**. A) Representative western blot after Co-IP for H2B and H2BE. B) Extracted ion chromatograms (EICs) for H2B peptides (top) and H2BE peptides (bottom), showing the successful differentiation between the two. C) Number of mapped reads of either H2BE (pink) or H2B (grey) with histone variants. D) Same as C, this time for H1 linker histones. E) GO biological process analysis of H2B-enriched binding partners.

**Supplementary Figure 2. The BAF complex promotes H2BE incorporation in chromatin.** A) Western blot for H2BE expression after 6 and 24 hours of 10uM BAFi. B) Metaplot of H2BE CUT&Tag profiling in WT primary cortical neurons treated with DMSO (grey) or BAFi (pink). The plot shows read counts per million mapped reads (RPM) at ± 2 kb from the transcription start site (TSS) of all genes expressed in mouse neurons. N = 3 biological replicates per treatment. C) Heatmap and metaplot of H2BE signal +/-1KB of the TSS across all genes. D) Normalized H2BE CUT&Tag read counts at H2BE peak sites with ≥ 10 reads detected at +/- 500bp of the TSS (unpaired t test). E) GO biological process analysis of lost H2BE peaks in BAFi condition using a background list of genes expressed in mouse neurons. F) Metaplot and heatmaps of H2BE signal +/-1KB of the TSS subset by gene expression. “Not expressed” was defined as genes with mean normalized read counts <3 by RNA-sequencing^15^. Remaining genes were binned into two equally sized groups by mean normalized read counts. G) Overlap of H2BE and H2A.Z peaks in the DMSO control condition (baseline). H) Overlap of H2BE and H2A.Z peaks lost with BAFi treatment. I) Log2fold change of H2A.Z and H2BE peaks with BAFi (unpaired t-test). J) qPCR assessing *Smarcd1* depletion in N2A cells (left) and neurons (right). Results were normalized to GAPDH. K) Western blots for SMARCD1 and H2BE expression after *Smarcd1* knockdown. L) Metaplot of H2BE CUT&Tag profiling in WT primary cortical neurons transduced with a Control shRNA (grey) or *Smarcd1* targeting shRNA (pink). N = 3 biological replicates per treatment. M) Heatmap and metaplot of H2BE signal +/- 1KB of the TSS across all genes. N) Normalized H2BE CUT&Tag read counts at H2BE peak sites with ≥ 10 reads detected +/- 500bp of the TSS (unpaired t test). O) GO biological process analysis of lost H2BE peaks with *Smarcd1* depletion using a background list of genes expressed in mouse neurons. P) Heatmap and metaplots +/- 500bp from the TSS, subset by gene expression. “Not expressed” was defined as genes with mean normalized read counts <3 by RNA-sequencing. Remaining genes were binned into two equally sized groups by mean normalized read counts. Q) Scatterplot correlation of BAFi and *Smarcd1* depletion H2BE CUT&Tag normalized counts. Pearson r = 0.80.

**Supplementary Figure 3. SP1 restricts BAF-mediated H2BE deposition.** A) CoIP in N2As for the association between SP1 and SMARCD1. D) Heatmap and metaplots +/-1KB from the TSS, subset by gene expression. “Not expressed” was defined as genes with mean normalized read counts <3 by RNA-sequencing^15^. Remaining genes were binned into two equally sized groups by mean normalized read counts. E) Metaplot of H2BE CUT&Tag profiling in WT primary cortical neurons treated with DMSO (grey) or Flavopiridol (light green). Plot shows read counts per million mapped reads (RPM) at ± 2 kb from the transcription start site (TSS) of genes expressed in mouse neurons. N = 3 biological replicates per treatment. F) Heatmap and metaplot of H2BE signal +/- 1KB of the TSS across all genes. G) Normalized H2BE CUT&Tag read counts at H2BE peak sites with ≥ 10 reads detected at the +/- 500bp of the TSS (unpaired t-test). H) Overlap of H2BE peaks in DMSO and Flavopiridol at the TSS called by SEACR. Log2fold change of H2BE (treatment – control) in Flavopiridol and *Sp1* depletion experiments. J) Overlap of gained H2BE peaks with *Sp1* depletion and Flavopiridol treatment.

**Supplementary Figure 4. The FACT complex restricts H2BE to the TSS.** A) CoIP in neurons for the association between H2BE and FACT subunit SSRP1. B) Same as A but in neurons. C) Western blot for H2BE and H2B after treatment with FACTi for 6, 24, and 46 hours. D) Metaplot and heatmap at +/- 1KB from the TSS of all genes for DMSO and FACTi. E) GO plot depicting gained H2BE genic peaks with FACTi treatment. F) Heatmap and metaplots +/- 1KB from the TSS, subset by gene expression. “Not expressed” was defined as genes with mean normalized read counts <3 by RNA-sequencing^17^. Remaining genes were binned into two equally sized groups by mean normalized read counts. G) Normalized H2BE CUT&Tag read counts at H2BE peak sites with ≥ 10 reads detected at genic regions minus first and last 500 bp, subset by gene expression (unpaired t test; ***, p < 0.001).

**Supplementary Figure 5. NAP1L4 evicts H2BE from chromatin.** A) qPCR of *Nap1l4* shRNA knockdown in N2As and neurons. Results were normalized to GAPDH. B) Western blot of *Nap1l4* knockdown in N2As and neurons. C) GO biological process analysis of genes that gained an H2BE peak in the *Nap1l4* knockdown using a background list of genes expressed in mouse neurons. D) Heatmap and metaplots +/-1KB from the TSS, subset by gene expression. “Not expressed” was defined as genes with mean normalized read counts <3 by RNA-sequencing^15^. Remaining genes were binned into two equally sized groups by mean normalized read counts. E) Overlap of H2BE peaks gained at the TSS in *Nap1l4* knockdown, compared to Flavopiridol treatment. F) Representative CUT&Tag gene tracks showing H2BE localization in *Nap1l4* knockdown and Flavopiridol conditions. Bigwigs were averaged across replicates.

**Supplementary Figure 6. H2BE-dependent transcriptional effects of chaperone depletion.** A) Volcano plot of FACTi vs DMSO in WT neurons, n = 4 biological replicates/treatment. Vertical lines represent a log2-fold change cutoff of 0.5 (absolute value) and horizontal lines represent a p-value cutoff of 0.001. B) Volcano plot of SP1i vs DMSO in WT neurons, n = 4 biological replicates/treatment. Vertical lines represent a log2-fold change cutoff of 0.5 (absolute value) and horizontal lines represent a p-value cutoff of 0.001. C) GO biological process analysis of genes that were downregulated with BAFi using a background list of genes expressed in mouse neurons. D) GO biological process analysis of genes that were upregulated with BAFi using a background list of genes expressed in mouse neurons. E) Representative gene track of a gene that lost H2BE at its TSS with BAF perturbation but had increased gene expression. For CUT&Tag, bigwigs were generated from merged replicates. For RNA-seq, tdfs were generated from individual replicates and then merged. F) Venn diagram of genes downregulated in BAFi vs DMSO in WT neurons vs genes downregulated in H2BE KO vs WT neurons. G) Venn diagram of genes upregulated in BAFi vs DMSO in WT neurons vs genes upregulated in H2BE KO vs WT neurons. H) Heatmap of z-scored CPM of a subset of genes that were shared in the comparison in Figure 6G. I) Log2fold changes (BAFi – DMSO) of genes that were downregulated both by H2BE KO compared to WT H2BE (Down, H2BE KO) and by BAFi in a WT context (Down, WT BAFi). *** p < 0.001 in an unpaired t-test. J) GO plot of genes that went up only in the WT *Nap1l4* depletion context, using a background list of genes expressed in mouse neurons. K) GO plot of genes that went down only in the WT *Nap1l4* depletion context, using a background list of genes expressed in mouse neurons. L) Heatmap of genes that went up in WT *Nap1l4* depletion but not in H2BE KO neurons. Clustering represents 3 major patterns identified. The dashed line represents the cluster from which Figure 6N was subset. M) H2BE-independent gene that was upregulated in both WT and KO *Nap1l4* depleted neurons. N) Example gene track of an H2BE-independent gene that is upregulated in WT neurons with *Nap1l4* depletion *and* upregulated in the Control H2BE KO condition.

