## Supplementary Figures for "Mechanisms controlling the deposition and dynamics of histone variant H2BE"

#### Supplementary Figure 1

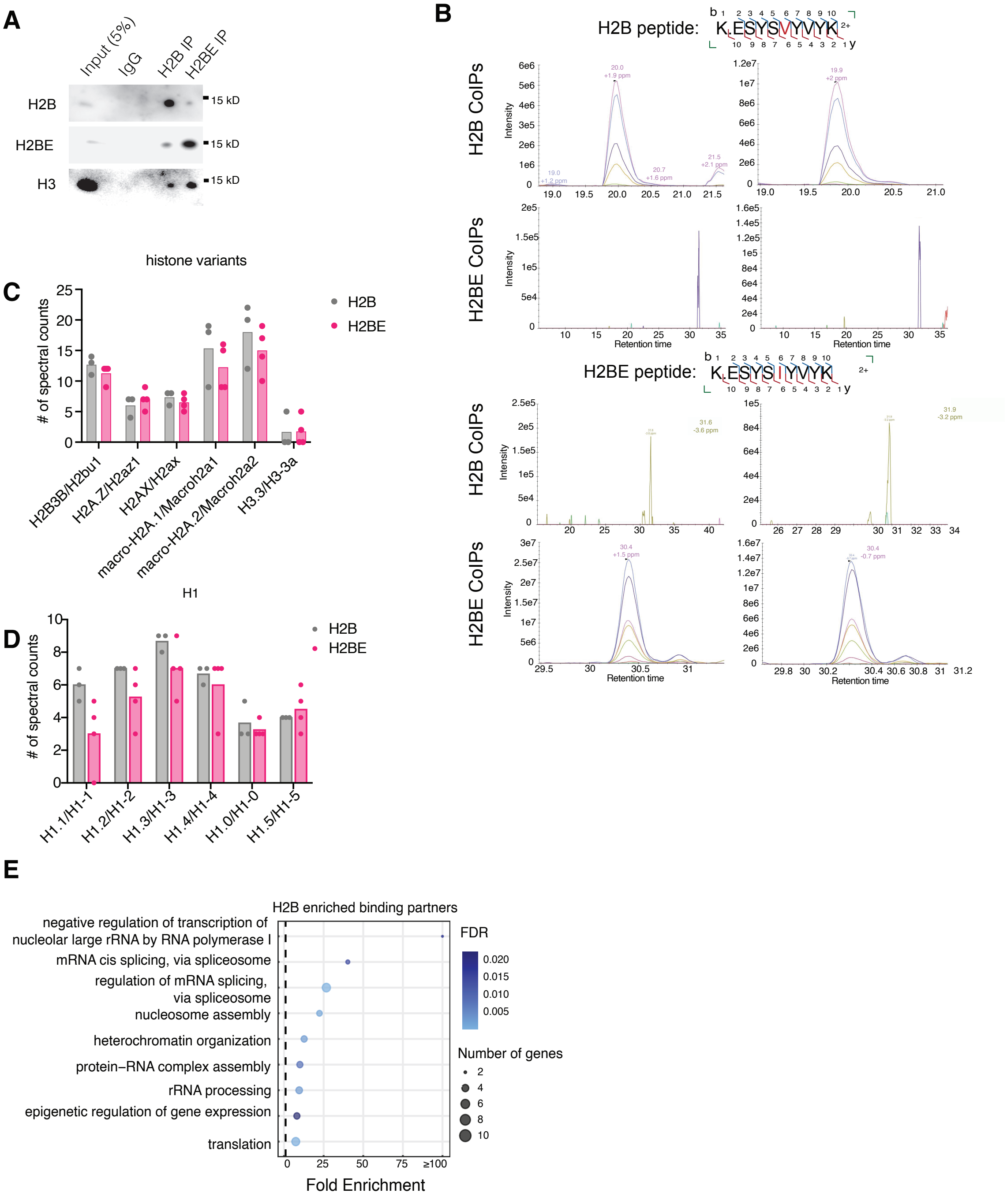


**Supplementary Figure 1. Identification of H2BE binding partners**. A) Representative western blot after Co-IP for H2B and H2BE. B) Extracted ion chromatograms (EICs) for H2B peptides (top) and H2BE peptides (bottom), showing the successful differentiation between the two. C) Number of mapped reads of either H2BE (pink) or H2B (grey) with histone variants. D) Same as C, this time for H1 linker histones. E) GO biological process analysis of H2B-enriched binding partners.

### Supplementary Figure 2

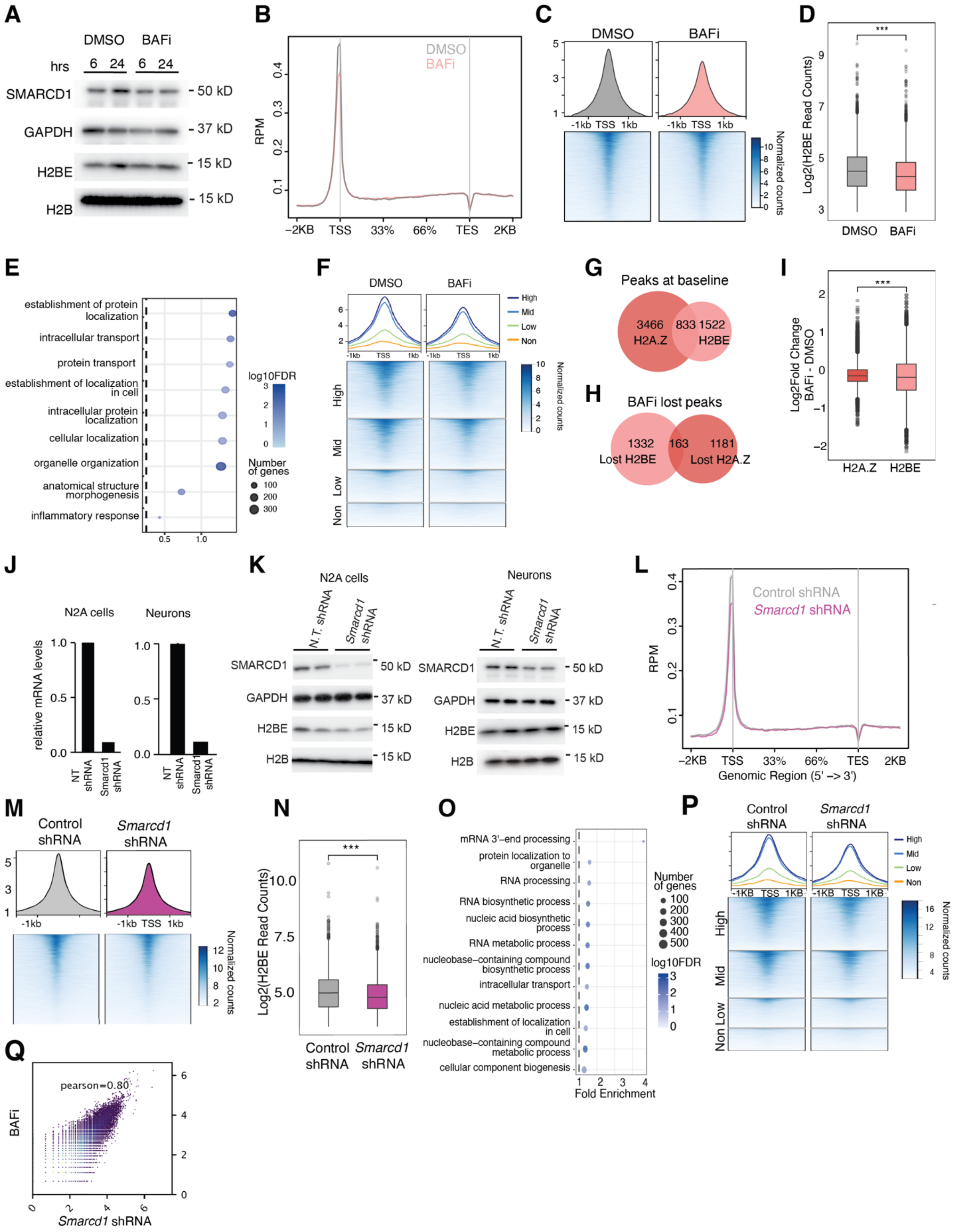

**Supplementary Figure 2. The BAF complex promotes H2BE incorporation in chromatin**. A) Western blot for H2BE expression after 6 and 24 hours of 10uM BAFi. B) Metaplot of H2BE CUT&Tag profiling in WT primary cortical neurons treated with DMSO (grey) or BAFi (pink). The plot shows read counts per million mapped reads (RPM) at ± 2 kb from the transcription start site (TSS) of all genes expressed in mouse neurons. N = 3 biological replicates per treatment. C) Heatmap and metaplot of H2BE signal +/- 1KB of the TSS across all genes. D) Normalized H2BE CUT&Tag read counts at H2BE peak sites with ≥ 10 reads detected at +/- 500bp of the TSS (unpaired t test). E) GO biological process analysis of lost H2BE peaks in BAFi condition using a background list of genes expressed in mouse neurons. F) Metaplot and heatmaps of H2BE signal +/- 1KB of the TSS subset by gene expression. “Not expressed” was defined as genes with mean normalized read counts <3 by RNA-sequencing[15]. Remaining genes were binned into two equally sized groups by mean normalized read counts. G) Overlap of H2BE and H2A.Z peaks in the DMSO control condition (baseline). H) Overlap of H2BE and H2A.Z peaks lost with BAFi treatment. I) Log2fold change of H2A.Z and H2BE peaks with BAFi (unpaired t-test). J) qPCR assessing *Smarcd1* depletion in N2A cells (left) and neurons (right). Results were normalized to GAPDH. K) Western blots for SMARCD1 and H2BE expression after *Smarcd1* knockdown. L) Metaplot of H2BE CUT&Tag profiling in WT primary cortical neurons transduced with a Control shRNA (grey) or *Smarcd1* targeting shRNA (pink). N = 3 biological replicates per treatment. M) Heatmap and metaplot of H2BE signal +/- 1KB of the TSS across all genes. N) Normalized H2BE CUT&Tag read counts at H2BE peak sites with ≥ 10 reads detected +/- 500bp of the TSS (unpaired t test). O) GO biological process analysis of lost H2BE peaks with *Smarcd1* depletion using a background list of genes expressed in mouse neurons. P) Heatmap and metaplots +/- 500bp from the TSS, subset by gene expression. “Not expressed” was defined as genes with mean normalized read counts <3 by RNA-sequencing. Remaining genes were binned into two equally sized groups by mean normalized read counts. Q) Scatterplot correlation of BAFi and *Smarcd1* depletion H2BE CUT&Tag normalized counts. Pearson r = 0.80.

#### Supplementary Figure 3

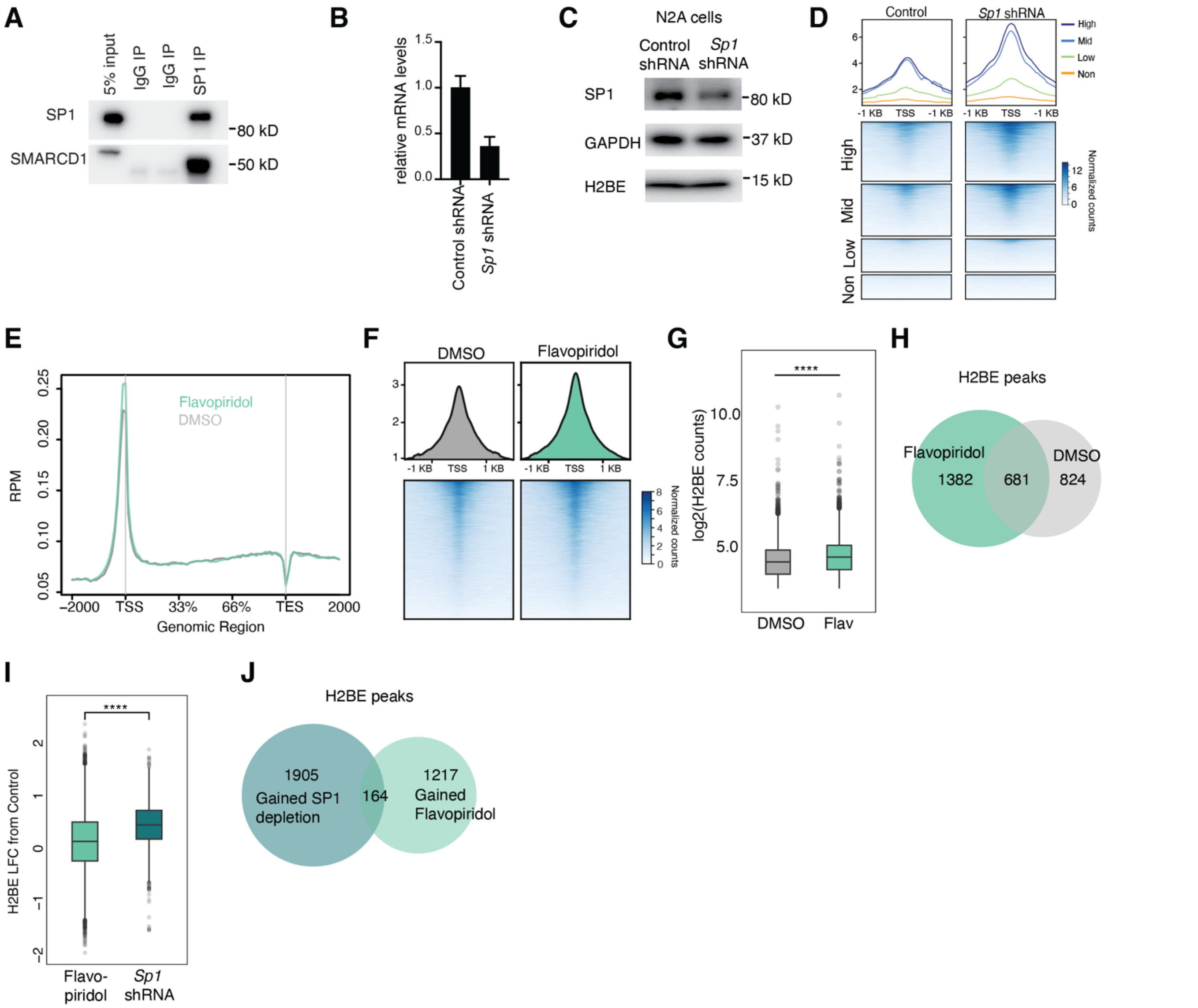


**Supplementary Figure 3. SP1 restricts BAF-mediated H2BE deposition.** A) CoIP in N2As for the association between SP1 and SMARCD1. D) Heatmap and metaplots +/- 1KB from the TSS, subset by gene expression. "Not expressed" was defined as genes with mean normalized read counts <3 by RNA-sequencing[15]. Remaining genes were binned into two equally sized groups by mean normalized read counts. E) Metaplot of H2BE CUT&Tag profiling in WT primary cortical neurons treated with DMSO (grey) or Flavopiridol (light green). Plot shows read counts per million mapped reads (RPM) at ± 2 kb from the transcription start site (TSS) of genes expressed in mouse neurons. N = 3 biological replicates per treatment. F) Heatmap and metaplot of H2BE signal +/- 1KB of the TSS across all genes. G) Normalized H2BE CUT&Tag read counts at H2BE peak sites with ≥ 10 reads detected at the +/- 500bp of the TSS (unpaired t-test). H) Overlap of H2BE peaks in DMSO and Flavopiridol at the TSS called by SEACR. Log2fold change of H2BE (treatment – control) in Flavopiridol and *Sp1* depletion experiments. J) Overlap of gained H2BE peaks with *Sp1* depletion and Flavopiridol treatment.

#### Supplementary Figure 4

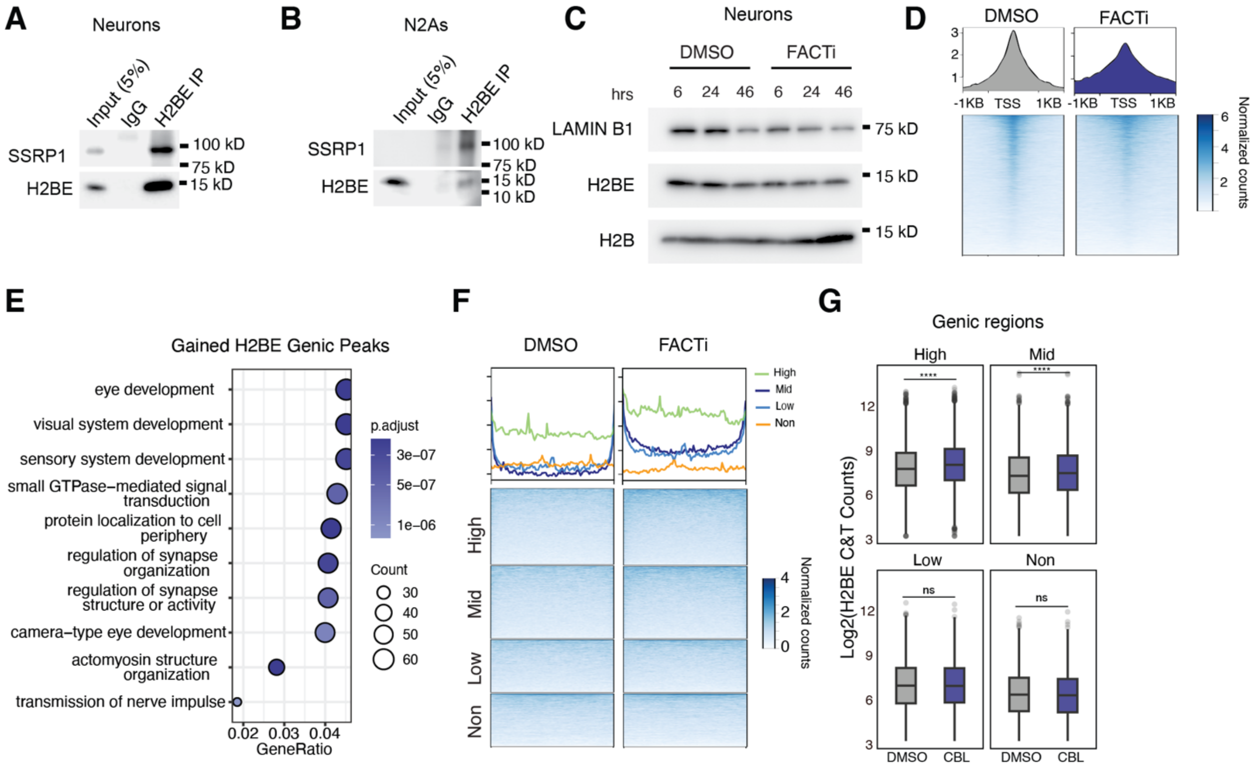


**Supplementary Figure 4. The FACT complex restricts H2BE to the TSS.** A) CoIP in neurons for the association between H2BE and FACT subunit SSRP1. B) Same as A but in neurons. C) Western blot for H2BE and H2B after treatment with FACTi for 6, 24, and 46 hours. D) Metaplot and heatmap at +/- 1KB from the TSS of all genes for DMSO and FACTi. E) GO plot depicting gained H2BE genic peaks with FACTi treatment. F) Heatmap and metaplots +/- 1KB from the TSS, subset by gene expression. “Not expressed” was defined as genes with mean normalized read counts <3 by RNA-sequencing[15]. Remaining genes were binned into two equally sized groups by mean normalized read counts. G) Normalized H2BE CUT&Tag read counts at H2BE peak sites with ≥ 10 reads detected at genic regions minus first and last 500 bp, subset by gene expression (unpaired t-test).

#### Supplementary Figure 5

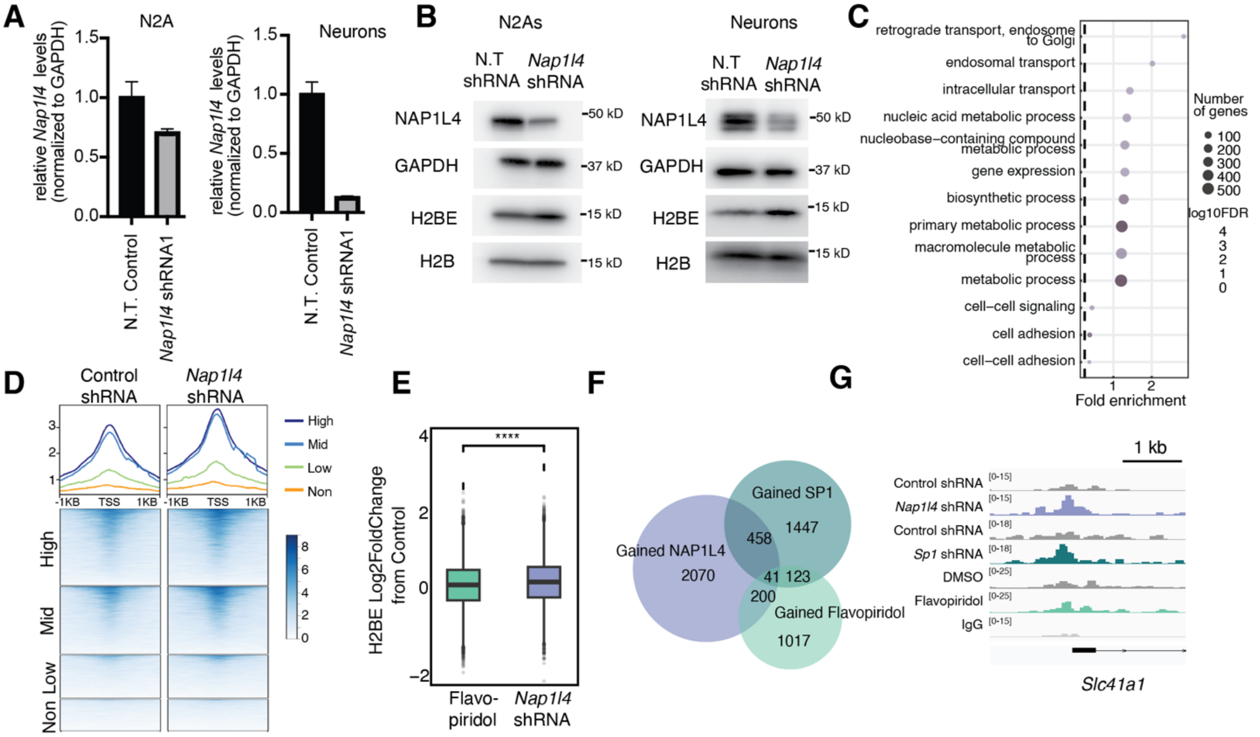

**Supplementary Figure 5. NAP1L4 evicts H2BE from chromatin**. A) qPCR of *Nap1l4* shRNA knockdown in N2As and neurons. Results were normalized to GAPDH. B) Western blot of *Nap1l4* knockdown in N2As and neurons. C) GO biological process analysis of genes that gained an H2BE peak in the *Nap1l4* knockdown using a background list of genes expressed in mouse neurons. D) Heatmap and metaplots +/- 1KB from the TSS, subset by gene expression. “Not expressed” was defined as genes with mean normalized read counts <3 by RNA-sequencing[15]. Remaining genes were binned into two equally sized groups by mean normalized read counts. E) Overlap of H2BE peaks gained at the TSS in *Nap1l4* knockdown, compared to Flavopiridol treatment. F) Representative CUT&Tag gene tracks showing H2BE localization in *Nap1l4* knockdown and Flavopiridol conditions. Bigwigs were averaged across replicates.

### Supplementary Figure 6

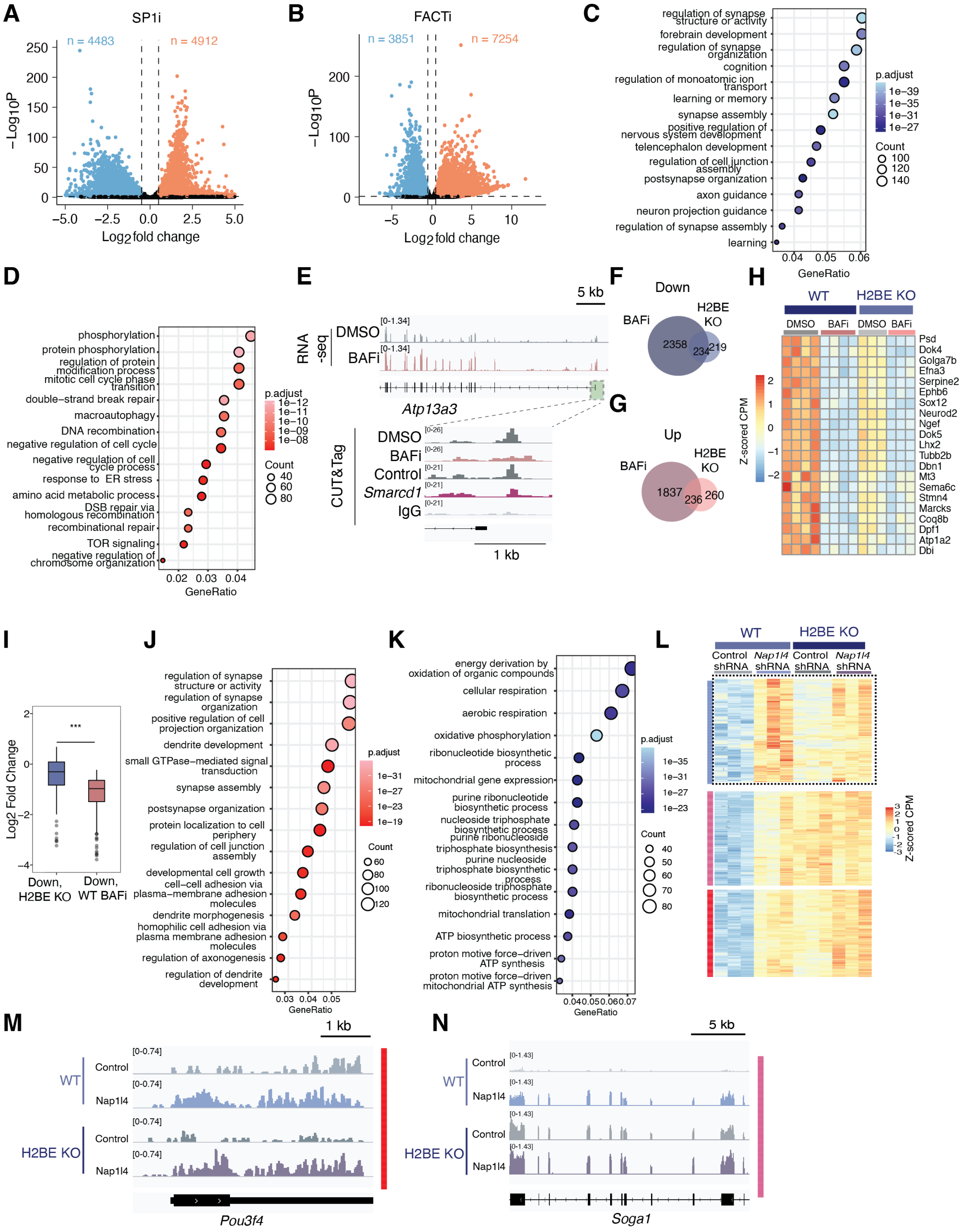

**Supplementary Figure 6**. **H2BE-dependent transcriptional effects of chaperone depletion**. A) Volcano plot of FACTi vs DMSO in WT neurons, n = 4 biological replicates/treatment. Vertical lines represent a log2-fold change cutoff of 0.5 (absolute value) and horizontal lines represent a p-value cutoff of 0.001. B) Volcano plot of SP1i vs DMSO in WT neurons, n = 4 biological replicates/treatment. Vertical lines represent a log2-fold change cutoff of 0.5 (absolute value) and horizontal lines represent a p-value cutoff of 0.001. C) GO biological process analysis of genes that were downregulated with BAFi using a background list of genes expressed in mouse neurons. D) GO biological process analysis of genes that were upregulated with BAFi using a background list of genes expressed in mouse neurons. E) Representative gene track of a gene that lost H2BE at its TSS with BAF perturbation but had increased gene expression. For CUT&Tag, bigwigs were generated from merged replicates. For RNA-seq, tdfs were generated from individual replicates and then merged. F) Venn diagram of genes downregulated in BAFi vs DMSO in WT neurons vs genes downregulated in H2BE KO vs WT neurons. G) Venn diagram of genes upregulated in BAFi vs DMSO in WT neurons vs genes upregulated in H2BE KO vs WT neurons. H) Heatmap of z-scored CPM of a subset of genes that were shared in the comparison in Figure 6G. I) Log2fold changes (BAFi – DMSO) of genes that were downregulated both by H2BE KO compared to WT H2BE (Down, H2BE KO) and by BAFi in a WT context (Down, WT BAFi). *** $p < 0.001$ in an unpaired t-test. J) GO plot of genes that went up only in the WT *Nap1l4* depletion context, using a background list of genes expressed in mouse neurons. K) GO plot of genes that went down only in the WT *Nap1l4* depletion context, using a background list of genes expressed in mouse neurons. L) Heatmap of genes that went up in WT *Nap1l4* depletion but not in H2BE KO neurons. Clustering represents 3 major patterns identified. The dashed line represents the cluster from which Figure 6N was subset. M) H2BE-independent gene that was upregulated in both WT and KO *Nap1l4* depleted neurons. N) Example gene track of an H2BE-independent gene that is upregulated in WT neurons with *Nap1l4* depletion *and* upregulated in the Control H2BE KO condition.
